# Mutant p63 affects epidermal cell identity through rewiring the enhancer landscape

**DOI:** 10.1101/387902

**Authors:** Jieqiong Qu, Sabine Tanis, Jos P.H. Smits, Evelyn N. Kouwenhoven, Martin Oti, Ellen H. van den Bogaard, Colin Logie, Hendrik G. Stunnenberg, Hans van Bokhoven, Klaas Mulder, Huiqing Zhou

**Affiliations:** Department of Molecular Developmental Biology, Faculty of Science, Radboud Institute for Molecular Life Sciences, Radboud University, Nijmegen, the Netherlands; Department of Dermatology, Radboud Institute for Molecular Life Sciences, Radboud university medical center, Nijmegen, the Netherlands; Department of Molecular Biology, Faculty of Science, Radboud Institute for Molecular Life Sciences, Radboud University, Nijmegen, the Netherlands; Department of Human Genetics, Radboud university medical center, Nijmegen, the Netherlands; Department of Cognitive Neurosciences, Donders Institute for Brain, Cognition and Behavior, Radboud university medical center, Nijmegen, The Netherlands

**Author notes:** Correspondence: Huiqing Zhou (;). These authors contribute to the manuscript equally (Sabine Tanis, Jos P.H. Smits).

**Keywords:** p63, enhancer, cell identity, transcriptional regulation, EEC syndrome

## Abstract

Transcription factor p63 is a key regulator of epidermal keratinocyte proliferation and differentiation. In humans mutations in p63 cause several developmental disorders with defects of ectoderm-derived structures including the epidermis. The underlying molecular mechanisms of these mutations however remain unclear. Here we characterized the transcriptome and epigenome from EEC syndrome patients carrying mutations in the p63 DNA-binding domain. The transcriptome of p63 mutant keratinocytes deviated from the normal epidermal cell identity. Epigenomic analyses showed that the deregulated gene expression in p63 mutant keratinocytes resulted from an altered enhancer landscape contributed by loss of p63-bound active enhancers and by unexpected gain of enhancers. The gained enhancers in mutant keratinocytes were frequently bound by deregulated transcription factors such as RUNX1. Reversing RUNX1 overexpression partially rescued deregulated gene expression as well as the enhancer distribution. Our findings support the pivotal role of p63 in controlling the enhancer landscape of epidermal keratinocytes and identify a novel mechanism whereby p63 DNA-binding mutations associated with EEC syndrome rewire the enhancer landscape and affect epidermal cell identity.

## Introduction

The transcription factor (TF) p63 is an ancient member of the p53 gene family. Different from p53 that has a convincing function in tumor suppression, p63 is a key regulator for the development of the epidermis, specifically in epidermal stem cell self-renewal, morphogenesis and directing differentiation programs (Candi, Cipollone et al., 2008, Mills, Zheng et al., 1999, Yang, Schweitzer et al., 1999). Several p63 isoforms have been reported including the N-terminal TA and ΔN isoforms, and multiple C-terminal isoforms (Kouwenhoven, van Bokhoven et al., 2015b). All isoforms contain the DNA-binding domain and the oligomerization domain responsible for tetramerization of p63 (Browne, Cipollone et al., 2011).

The role of p63 in epidermal development has been established by two independent p63 knockout mouse models (Mills et al., 1999, Yang et al., 1999). These p63-deficient mice do not have stratified epidermis and epidermal related appendages. During embryonic development, p63-deficient mice develop a normal ectoderm with Krt8/Krt18 positive simple epithelial cells. However, they fail to initiate embryonic stratification and to produce mature Krt5/Krt14 positive epithelial and epidermal cells, termed as keratinocytes (Shalom-Feuerstein, Lena et al., 2011). Furthermore, p63-deficiency results in the activation of mesodermal genes during development (Shalom-Feuerstein et al., 2011). These findings demonstrate that p63 is essential and required for the commitment to a proper epidermal cell fate during development.

In keratinocytes, p63 plays important roles for both proliferation and differentiation. The p63 protein, mainly the ΔNp63α isoform, is expressed at a high level in proliferating keratinocytes in the basal layer of the epidermis. Upon stratification, its expression level is reduced (Candi, Dinsdale et al., 2007). Knockdown of p63 in keratinocytes affects proliferation and prevents cells from differentiation (Truong, Kretz et al., 2006). At the molecular level, knockdown of p63 induces genes controlling cell cycle arrest such as p21 (*CDKN1A*) (LeBoeuf, Terrell et al., 2010) and downregulates genes that are important for epidermal differentiation such as *PERP* and *KRT14* (Ihrie, Marques et al., 2005, Romano, Birkaya et al., 2007). Many of these genes are direct p63 target genes. These data show that p63 represses cell cycle arrest genes to promote proliferation, and activates epidermal differentiation genes to induce differentiation.

In recent years, a number of epigenomic profiling studies firmly established the master regulator role of p63 in the genome of keratinocytes, predominantly in controlling enhancers (Cavazza, Miccio et al., 2016, Kouwenhoven, Oti et al., 2015a, Kouwenhoven et al., 2015b, Lin-Shiao, Lan et al., 2018, Rinaldi, Datta et al., 2016). p63 bookmarks genomic loci, and cooperates with specific TFs to activate epidermal genes via active enhancers. Consistently, ATAC-seq analysis showed that p63 binding sites are preferentially located in nucleosome-enriched regions in epidermal keratinocytes, and these sites are inaccessible in cell types where p63 is not expressed (Bao, Rubin et al., 2015). In keratinocytes, p63 cooperates with an ATP-dependent chromatin remodeling factor BAF1 to make these regions accessible. It has also been shown that p63 directly regulates chromatin factors such as Satb1 and Brg1 that play roles in higher-order chromatin remodeling, covalent histone modifications, and nuclear assembly (Fessing, Mardaryev et al., 2011, Mardaryev, Gdula et al., 2014). These data suggest that p63 regulates epidermal cell fate determination and differentiation not only through direct target genes but also via modulating the epigenetic landscape.

The key role of p63 in epidermal development has also been demonstrated in human disease models. Heterozygous mutations of *TP63* encoding p63 cause a spectrum of developmental disorders. Among them, Ectrodactyly-Ectodermal Dysplasia-Cleft Lip/Palate (EEC) syndrome is caused by point mutations located in the p63 DNA-binding domain, and manifests ectodermal dysplasia with defects in the epidermis and epidermal related appendages, limb malformation and cleft lip/palate (Rinne, Brunner et al., 2007). Five hotspot mutations affecting amino acids, R204, R227, R279, R280 and R304, have been found in approximately 90% of the EEC population, and these EEC mutations were shown to disrupt p63 DNA binding and result in impaired transactivation activity (Browne et al., 2011, Brunner, Hamel et al., 2002, Celli, Duijf et al., 1999, Rinne et al., 2007). Therefore, these mutant p63 proteins have been proposed to have a dominant negative effect towards the wild type p63, probably by abolishing DNA binding as a result of tetramerization of the wild type and mutant proteins (Brunner et al., 2002). Furthermore, mouse genetic studies support the dominant negative model. Heterozygous p63 knockout mice do not show any ectodermal phenotype (Mills et al., 1999, Yang et al., 1999), whereas heterozygous knockin mice carrying an EEC mutation resemble the human phenotype (Vernersson Lindahl, Garcia et al., 2013).

Although the role of p63 in normal epidermal development and differentiation has been demonstrated, the molecular mechanism by which p63 mutations cause the epidermal phenotype in diseases is not yet understood. We previously reported that p63 mutant keratinocytes derived from EEC patients could not fully differentiate towards terminal stratification in both 2D and 3D cellular models (Shen, van den Bogaard et al., 2013). In this study, EEC patient keratinocytes carrying three hotspot mutations, R204W, R279H and R304W, were assessed by transcriptomic and epigenomic analyses to identify the underlying molecular mechanism. Our data showed that deregulated gene expression accompanied by a rewired enhancer landscape led to a less defined epidermal cell identity of p63 mutant keratinocytes, which potentially contributes to the pathogenic mechanism of EEC syndrome.

## Results

### Loss of characteristic epidermal expression profiles in p63 mutant keratinocytes

Using an established *in vitro* differentiation model of epidermal keratinocytes (Kouwenhoven et al., 2015a), we characterized gene expression differences between keratinocytes derived from non-EEC individuals (control) and from EEC patients carrying mutations in the DNA-binding domain of p63 (R204W, R279H and R304W, p63 mutants) (Figs 1A and B). Similar to our previous reports (Shen et al., 2013), p63 mutant keratinocytes remained largely unchanged morphologically at the terminal stage of differentiation, as compared to the multilayer cell structures of control keratinocytes on day 7 (Fig EV1A), indicating that they were unable to fully differentiate. To better characterize these mutant keratinocytes at the molecular level, we performed paired-end RNA-seq analyses. In Principal Component Analysis (PCA), gene expression of control keratinocytes from day 0 to day 7 moved along the PC1 axis (51%) that probably defines the differentiation process, whereas mutant keratinocytes remained at the left side of PC1 (Fig 1C). Consistently, DAVID Gene Ontology (GO) annotation (Dennis, Sherman et al., 2003) of the top 500 genes with positive weights in PC1 showed terms of ‘epidermis development’ and ‘keratinocyte differentiation’ (Tables EV1A and B). Many affected epidermal genes including epidermal marker genes, *KRT5, KRT1, KRT10*, and genes from the epidermal differentiation complex (EDC), e.g. *involucrin (IVL), filaggrin (FLG), FLG2, LCE2A, LCE5A*, and *loricrin (LOR)*, were validated by RT-qPCR (Fig EV1B). Furthermore, epidermal marker gene, *LOR* that was down-regulated at the end of differentiation, were also validated at the protein level (Fig EV1B). These molecular data confirmed the differentiation defect of p63 mutant keratinocytes shown in morphology.

**Figure 1.**
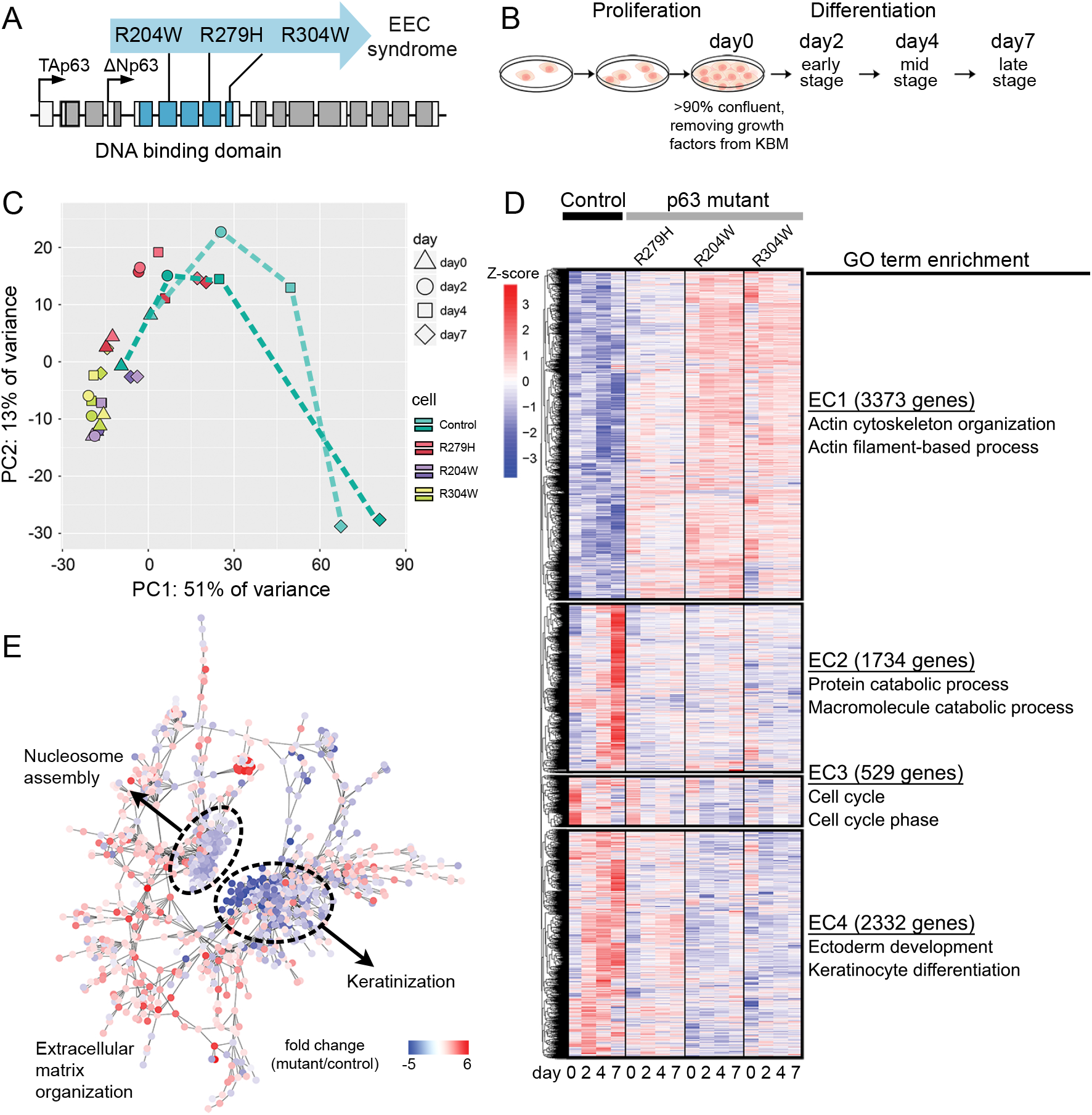
Transcriptome dynamics of control and p63 mutant keratinocytes during differentiation. A. Three EEC syndrome patient-derived primary keratinocyte lines, each carrying one mutation in the DNA binding domain (blue) of the *TP63* gene (p63 mutant keratinocytes). B. The setup of *in vitro* differentiation of epidermal keratinocytes. Samples were collected at four key stages, proliferation (day 0), early differentiation (day 2), mid differentiation (day 4), and late differentiation (day 7). C. Principal component analysis (PCA) on RNA-seq data. Two independent control lines (as biological replicates) and three p63 mutant lines are indicated with different colors. Shapes indicate four stages. D. Hierarchical clustering of differentially expressed genes (*P* value <0.05). Z-score was calculated based on log10 (FPKM+0.01) of each gene. Enriched top two Gene Ontology (GO) terms of genes per gene expression cluster (EC) are shown and all significant GO terms are summarized in Table EV1. E. Co-expression network of deregulated genes in p63 mutant keratinocytes during differentiation. Interactions with connectivity weight > 0.1 were shown. Two main co-expression modules were labelled with corresponding GO terms. The node color indicates the expression fold change between mutant and control keratinocytes. See also Fig EV1, Fig EV2 and Table EV1.

We next analyzed differentially expressed (DE) genes (*P* value < 0.05) between control and p63 mutant keratinocytes during differentiation. Overall, 3373 genes were upregulated and 4595 genes were downregulated in p63 mutant keratinocytes (Table EV1D), which were distinguished into four clusters by k-means clustering (Fig 1D and Table EV1E). Among the upregulated genes (EC1), some were generally not expressed or expressed at a low level in control keratinocytes. There was an enrichment of genes involved in extracellular structure organization, actin cytoskeleton organization (*LIMA1, TLN1*, and *PDGFB*) and muscle cell function (*CAV2, TNC, TTN*, and *MYH10*) (Table EV1F), which are consistent with previous reports that p63 deficiency resulted in upregulation of mesodermal genes (Barton, Tahinci et al., 2009, Shalom-Feuerstein et al., 2011). TFs in this cluster include *SOX4, TEAD2, RUNX1* that are widely expressed in many cell types and a number of Antp homeobox family members such as *HOX* genes, e.g. *HOXB3* and *HOXB8*. Interestingly, *TP63* was also detected in EC1, and ΔNp63 was the isoform detected in both control and mutant keratinocytes (Fig EV1C). Compared to the decreased p63 expression during differentiation in control keratinocytes, an increased p63 expression at the proliferation stage was observed in p63 mutant keratinocytes, which stayed at a high level through differentiation (Fig EV1C).

Expression of the 4595 genes whose expression was dynamically induced during differentiation in control keratinocytes remained low and largely unchanged in p63 mutant keratinocytes. These genes were grouped to three clusters (EC2, EC3, and EC4) (Tables EV1G-I). Genes in EC2 showed a sharp increase in gene expression on day 7 in control keratinocytes. Many of these genes were involved in catabolic pathways and cell death (e.g., *UBE4A* and *CDKN2D*). Genes in EC3 were highly expressed at the proliferation stage on day 0 and expression went down during differentiation. They were mainly involved in cell cycle regulation (e.g., *CDC20* and *KIFC1*). Finally, genes in EC4 showed a progressive upregulation in control keratinocytes and many of them were involved in ectoderm development and keratinocyte differentiation. The TF genes in this cluster included *RORA, OVOL1* and *KLF4*, and known p63 co-regulators, such as *MAFB, TFAP2A* and *NFE2L2* (Kouwenhoven et al., 2015a, McDade, Henry et al., 2012). In addition, many cell cycle related genes were affected, consolidating the role of p63 in cell cycle control. We validated some of these deregulated genes, e.g. *CDKN1A, CDKN2B*, and *MKI67* by RT-qPCR (Fig EV1B). Of note, consistent with the morphological changes (Fig EV1A), the deregulated gene expression was more evident in mutant keratinocytes carrying R204W and R304W than those carrying R279H (Fig 1D). Taken together, our RNA-seq analyses showed downregulation of epidermal differentiation genes and upregulation of non-epidermal genes in p63 mutant keratinocytes, suggesting that p63 mutant keratinocytes have a less defined epidermal cell identity.

To better visualize the interaction between DE genes, we carried out Weighted Gene-Coexpression Correlation Network Analyses using the Cytoscape Network Analyzer (http://www.cytoscape.org) (Tables EV1J and K). Two significant co-expression gene modules were identified, of which many genes were involved in ‘keratinization’ (e.g., *LOR, FLG* and *LCE3E*) and ‘nucleosome assembly’ (e.g., *HMGB2* and genes encoding histones) (Fig 1E, Fig EV2A and Table EV1L). Most genes in both modules showed downregulated expression in p63 mutant keratinocytes. The upregulated genes in p63 mutant keratinocytes did not generate significant main modules but several small sub-network modules. They likely played roles in ‘extracellular matrix organization’ (Table EV1L). However, the higher inter-modular connectivity between ‘keratinization’, ‘nucleosome assembly’ and ‘extracellular matrix organization’ modules suggests a biological relationship between these modules, indicating that changes in the chromatin landscape may contribute to gene deregulation. Consistent with this notion, many chromatin regulators were deregulated in p63 mutant keratinocytes (Fig EV2B). These factors include *KAT2B* which is a histone acetyltransferase (Marmorstein, 2001) and *SMYD3* which encodes a histone methyltransferase and functions in RNA polymerase II complexes (Hamamoto, Furukawa et al., 2004). The deregulation of these genes was confirmed with RT-qPCR (Fig EV2C).

### p63 orchestrates enhancer dynamics during epidermal differentiation

Given the indicated relationship between p63 and the chromatin landscape, we first assessed the role of p63 in regulating the epigenetic landscape during normal epidermal differentiation. We mapped histone modifications H3K27ac, H3K4me3, and H3K27me3, as well as p63 binding sites (BSs) of control keratinocytes (Kouwenhoven et al., 2015a) to open chromatin regions detected by DNase I Hypersensitivity Sites (DHSs) in Normal Human Epidermal Keratinocytes (NHEK) reported by ENCODE (Table EV2) (Myers, Stamatoyannopoulos et al., 2011). Two clusters of active enhancers (C3 and C4) were bound by p63 (Figs EV3A-C). Regions in C3 showed higher p63 binding signals. GO annotation using the Genomic Regions Enrichment of Annotation Tool (GREAT) which permits functional interpretation of *cis*-regulatory regions (McLean, Bristor et al., 2010) showed that nearby genes were involved in ‘apoptosis’ and ‘epidermis development’. Regions in C4 had relatively lower p63 binding signals and nearby genes were involved in ‘keratinocyte differentiation’ (Fig EV3C). Furthermore, cluster C7 represents a small group of open chromatin regions that were occupied by H3K27me3 signals and were devoid of p63 binding (Fig EV3A). Genes near these regions were associated with ‘pattern specification process’, such as ‘neuron fate commitment’ (Fig EV3C).

To quantify epigenetic dynamics, we used the ChromHMM Hidden Markov Model approach (Ernst & Kellis, 2012) to analyze the chromatin state transitions. With the combination of H3K27ac, H3K4me3 and H3K27me3, we obtained six classes of chromatin states, active enhancers, active promoters, unmodified regions, weak promoters, heterochromatin regions, and bivalent promoters (Fig 2A, Tables EV3A and B). By pair-wise comparison between two adjacent stages of differentiation, we observed major transitions between active enhancers and unmodified regions as well as between unmodified regions and heterochromatin regions (Fig 2B). As expected, we found that the transition from unmodified regions to active enhancers was generally associated with gene upregulation; *vice versa*, the transition from active enhancers to unmodified regions was associated with gene downregulation, at least at early differentiation stages (day 0 to day 2, and day 2 to day 4) (Fig 2C), e.g. regulation of *LOR* (Fig 2D) and *KRT1* (Fig EV3D). Furthermore, there were also transitions between active promoters and unmodified regions (Fig EV3E). Interestingly, many genes that are known to be expressed in cells of the mesodermal origin, e.g. *PAX2* (Bouchard, Souabni et al., 2002), were heavily marked by H3K27me3 in proliferating keratinocytes (day 0). The repression was relieved at the end of the terminal differentiation of keratinocytes (Fig EV3F).

**Figure 2.**
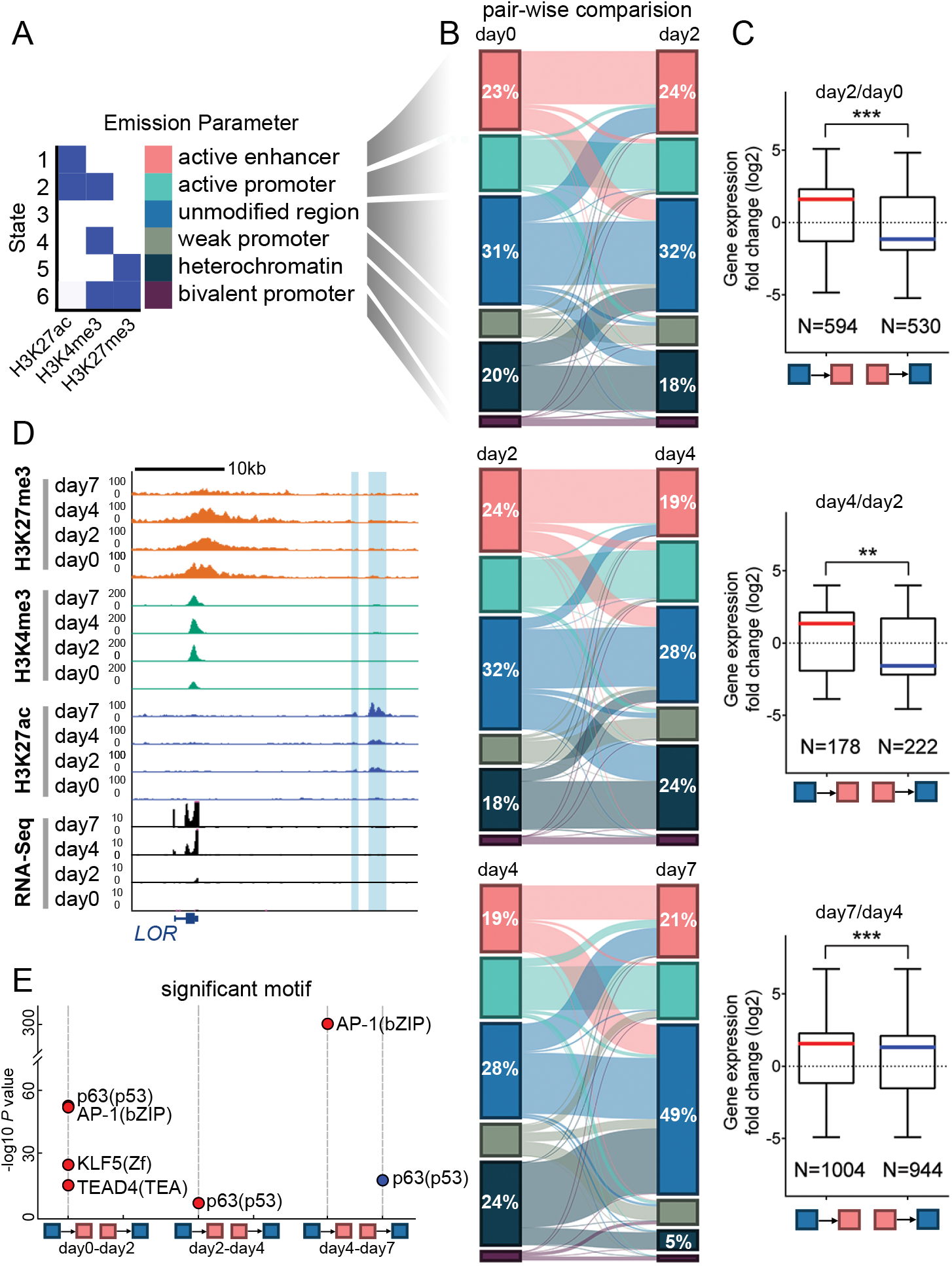
Chromatin dynamics during keratinocyte differentiation. A. Emission states of the ChromHMM Hidden Markov Model distinguishing six combinations defined by three histone modifications: H3K27ac, H3K4me3, and H3K27me3. ‘Unmodified regions’ refers to genomic regions that do not have any of these modifications. B. Alluvial plots of pair-wise chromatin state transitions during differentiation. The column height represents the percentage of each of the chromatin state relative to the sum of the six states. Genomic regions defined as ‘unmodified regions’ at all differentiation stages were excluded in this analysis. The percentages of active enhancers, unmodified regions, and the heterochromatin were labelled. C. Pair-wise comparison of differential gene expression (fold change) associated with genomic regions that shift between unmodified regions and active enhancers during differentiation, corresponding to B. Data are shown as mean ± standard deviation, unpaired T-test, ** *P* value <0.01, *** *P* value <0.001. D. Chromatin state dynamics of unmodified regions and active enhancers during differentiation at the locus of *LOR*. E. Significantly enriched motifs in the dynamic enhancer regions. Red dots, motifs that were enriched in regions shifting from unmodified regions to active enhancers; blue dots, motifs that were enriched in regions shifting from active enhancers to unmodified regions. See also Fig EV3, Tables EV2 and 3.

Next, we asked whether specific TFs control the enhancer dynamics during differentiation. Therefore, we performed motif analysis of dynamically changed enhancers with the HOMER package (http://homer.salk.edu/homer/motif/) (Fig 2E). We observed that bZIP, p53/p63, Zf, and TEA motifs were enriched in regions being activated from unmodified regions on day 0 to active enhancers on day 2, while p53/p63 was the only enriched motif in regions being activated from unmodified regions on day 2 to active enhancers on day 4. The bZIP motif was predominantly enriched in regions being activated from unmodified regions on day 4 to active enhancers on day 7, whereas the p53/p63 motif was the only enriched motif in regions changing from active enhancers on day 4 to unmodified regions on day 7 (Fig 2E). The temporal enrichment of p63 motifs in dynamic enhancers underscores the key role of p63 in orchestrating the enhancer landscape during keratinocyte differentiation.

### Decreased active enhancers associated with p63 binding deficiency in p63 mutant keratinocytes

Based on DNA-binding deficiency of EEC mutants and the dominant negative model shown by previous studies (Celli et al., 1999), we expected to detect DNA binding loss in p63 mutant keratinocytes. To characterize this and to evaluate the effect of p63 mutations on the enhancer landscape, we performed p63 ChIP-seq, using a p63 antibody that is not affected by mutations in the p63 DNA-binding domain (Shen et al., 2013), and H3K27ac ChIP-seq in all three p63 mutant keratinocytes. A total number of 33,366 p63 BSs detected in both control and mutant keratinocytes were analyzed, and we observed globally reduced p63 binding signals in p63 mutant keratinocytes as compared to the control keratinocytes (Fig 3A and Tables EV4A-E). It should be noted that no clear increased p63 binding or *de novo* p63 BSs were observed in p63 mutant keratinocytes, as compared to the control keratinocytes. K-means clustering analysis showed that p63 BSs can be clustered into three groups based on the binding signals. Clusters p63-C1 and p63-C2 had decreased p63 binding to a less extent, whereas loci in p63-C3 showed more dramatic to almost complete loss of p63 binding signals (Fig 3A). Accordingly, the difference of H3K27ac signals at p63-C1 and p63-C2 between the control and p63 mutant keratinocytes was not obvious, whereas a decrease was detected at p63 C3 in p63 mutant keratinocytes (Fig 3A and Fig EV4A).

**Figure 3.**
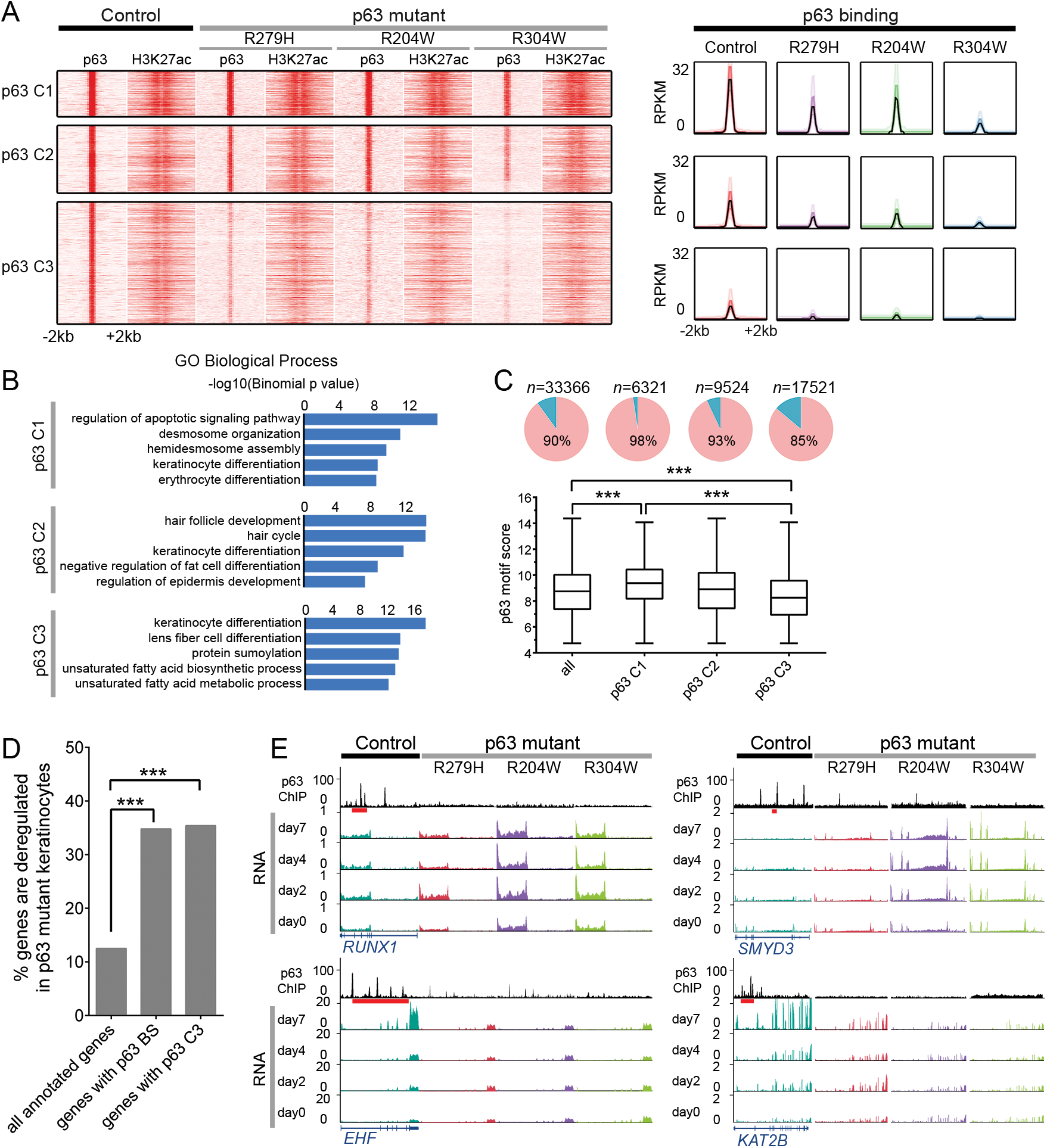
Decreased active enhancers associated with p63 binding deficiency in p63 mutant keratinocytes. A. K-means clustering of p63 binding sites (BSs) groups p63 BSs into 3 classes (*k* = 3, metric = Pearson). Merged p63 BSs from control and p63 mutant keratinocytes are shown in heat maps and band plots in a 4-kb window with summits of p63 BSs in the middle. Color intensity in heat maps represents normalized read counts. In the band plots, the median enrichment was visualized as the black line while 50% and 90% ranges were depicted in lighter color, respectively. B. GREAT-based Gene Ontology (GO) biological process annotation of p63 BSs in each cluster. C. p63 motif strength determined the selectivity of the loss of p63 binding. The top panel, pie charts showing the percentage of p63 BSs with a p63 motif; the bottom panel, box plot showing motif score distribution in all or different clusters of p63 BSs. Data are shown as mean ± standard deviation, one-way ANOVA, \*\*\**P* value < 0.001. D. Percentage of deregulated genes associated with all p63 BSs and p63 BSs from p63-C3 (p63 C3) compared with all annotated genes. Hypergeometric test, \*\*\**P* value < 0.001. E. ChIP-seq of p63 and RNA-seq data of *RUNX1, EHF, SMYD3* and *KAT2B* in control and p63 mutant keratinocytes. Red bars represent p63 BSs that were lost/decreased in p63 mutant keratinocytes. See also Fig EV4 and Table EV4.

Using GREAT GO annotation to assess nearby genes, all three clusters of p63 BSs were significantly enriched for genes involved in ‘epidermis development’ (Fig 3B). Interestingly, p63-C1 sites seemed to be associated more significantly to apoptosis and adhesion genes, whereas p63-C2 and p63-C3 sites to genes involved in ectodermal and epidermal function and structures such as hair follicles and keratinocytes (Fig 3B). We also performed human phenotype analyses using GREAT to investigate the disease significance of these p63 BSs, and detected disease terms were mainly related to ectodermal dysplasia such as ‘plantar hyperkeratosis’, ‘nail dystrophy’ and ‘alopecia’ (Fig EV4B).

We further explored whether a specific molecular mechanism controls the discordant p63 binding loss. We examined whether the cooperation with different co-regulating TFs contributes to the differential p63 binding loss. Motif scanning revealed that, in addition to p63/p53, bZIP, ETS, TEA and MAD motifs were enriched in all three p63 binding clusters, as compared to the genomic background (Tables EV4H-J), and therefore no significant differential p63 co-regulators were identified for the discordant p63 binding loss. Next, using our previously established p63scan algorithm (Kouwenhoven, van Heeringen et al., 2010), we found that p63-C3 BSs had the lowest percentage of BSs with p63 motif (85%) and the lowest average motif score (mean motif score of 8.3), as compared to p63-C1 (98% and mean motif score of 9.5) and p63-C2 (93% and mean motif score of 8.8) BSs (Fig 3D and Tables EV4K-N). Our observations thus suggest that p63 motif strength determines the selectivity of the loss of p63 binding.

Lastly, we examined whether gene deregulation was associated with impaired p63 binding. Genes near p63 BSs (all three p63-C1, C2 and C3) had a significantly larger proportion of deregulated genes (34.8%, *P* value = 0, hypergeometric test) when compared to all annotated genes (12.5%) (Fig 3D and Table EV4F). A significant difference in the percentage of deregulated genes was also found from genes associated with p63-C3 BSs (35.4%, *P* value = 0, hypergeometric test), as compared to all genes with p63 BSs (Tables EV4G). These data showed that impaired p63 binding significantly contributed to deregulated gene expression, both up- (e.g. *RUNX1* and *SMYD3)* and down-regulation (e.g. *EHF* and *KAT2B*) (Fig 3E).

In summary, we showed that EEC mutations can result in loss of p63 binding and loss of active enhancers. The loss of p63 binding is apparently motif strength-dependent. p63 binding loss can lead to gene deregulation and potentially contribute to ectodermal dysplasia phenotypes.

### Re-distribution of enhancers in p63 mutant keratinocytes

Although a significant percentage of the deregulated genes (~54%) associated with impaired p63 binding nearby, a large number of deregulated genes did not seem to be directly regulated by p63. Furthermore, we observed many enhancers with unexpected increased H3K27ac signals near deregulated genes in p63 mutant keratinocytes (Fig 4A, blue shaded regions). To further explore the mechanism, we performed comparison of H3K27ac between control and p63 mutant keratinocytes with MAnorm (Shao, Zhang et al., 2012). We identified 17,931 genomic regions that had significantly higher H3K27ac signals in mutant keratinocytes (referred to as ‘mutant-specific enhancers’), more than the 15,057 regions that had significantly higher H3K27ac in control keratinocytes (referred to as ‘control-specific enhancers’) (Fig 4B, Fig EV5A and Table EV5). It should be noted that, although these 17,931 ‘mutant-specific’ enhancers and 15,057 ‘control-specific enhancers’ showed significant quantitative difference in their H3K27ac signals, in most cases they were present in both control and mutant keratinocytes, rather than present exclusively in mutant or control cells, respectively. Two replicas of H3K27ac ChIP-seq in R304W mutant keratinocytes showed a high correlation (a Pearson correlation coefficient of 0.94, Fig EV5B), demonstrating the reliability of these analyses. To validate these findings, H3K27ac ChIP-qPCR was performed on three ‘control-specific’ and three ‘mutant-specific’ enhancer loci (Fig EV5C).

**Figure 4.**
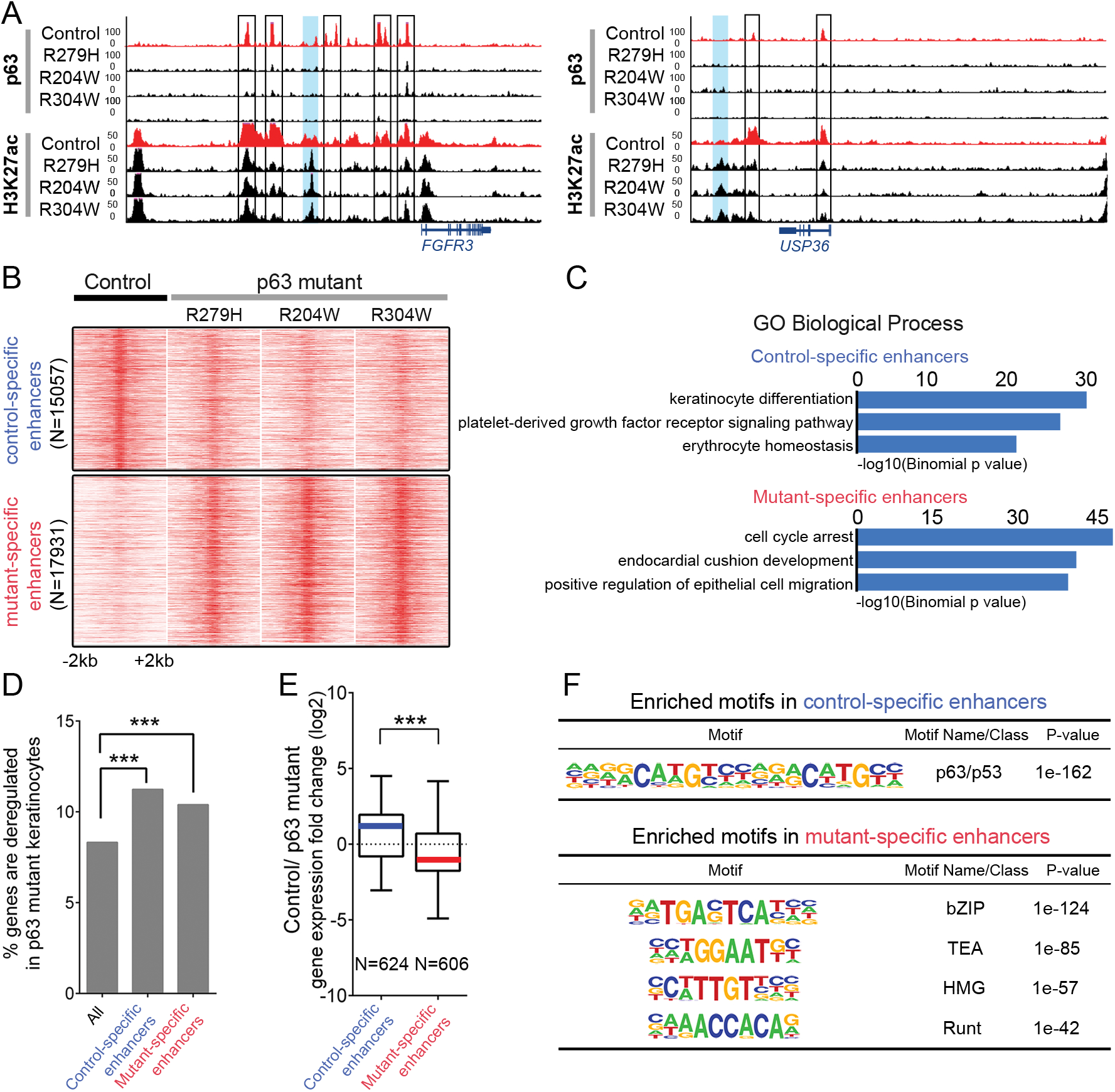
Redistribution of enhancers in p63 mutant keratinocytes. A. Examples of re-distributed enhancers marked by H3K27ac signals between control and p63 mutant keratinocytes. Black boxes indicate lost p63 BS and lost H3K27ac signals in p63 mutant keratinocytes at p63-bound enhancers detected in control keratinocytes. Blue shades indicate gained enhancers in p63 mutant keratinocytes. B. Heatmaps of differential H3K27ac peaks in control and p63 mutant keratinocytes shown in a 4-kb window with summits of H3K27ac peaks in the middle. Color intensity in heatmaps represents normalized read counts. C. GREAT-based GO biological process annotation of differential enhancers. D. Percentage of deregulated genes relative to all genes associated with active enhancers, all or control- or mutant-specific enhancers. Hypergeometric test, \*\*\**P* value < 0.001. E. Differential gene expression (fold change) associated with control or mutant-specific enhancers. Data are shown as mean ± standard deviation, unpaired T-test, *** *P* value <0.001. F. Highly enriched motifs found in control-specific or mutant-specific enhancers. See also Fig EV5, Tables EV5 and 6.

Genes nearby control-specific enhancers were associated with ‘keratinocyte differentiation’, whereas those nearby mutant-specific enhancers were involved in cell cycle regulation, migration and non-epithelial processes (Fig 4B), showing consistency with deregulated gene expression. Furthermore, a significantly larger proportion of deregulated genes had either control-specific or mutant-specific H3K27ac sites (Fig 4C). As expected, control-specific enhancers were associated with gene downregulation, whereas mutant-specific enhancers were associated with gene upregulation in p63 mutant keratinocytes when compared to control keratinocytes (Fig 4D). Among the 17,931 mutant-specific enhancers, only 5620 regions (31%) were not overlapping with DHSs in NHEK (ENCODE data). Interestingly, 2256/5620 regions (40%) were DHSs in HSMM (skeletal muscle myoblasts) and 2114/5620 regions (37%) were DHSs in NH-A (astrocytes). These data suggest that those ‘mutant-specific’ enhancers are potentially functional in other lineages, such as muscle cells or astrocytes. Taken together, the observed genome-wide redistribution of enhancers marked by H3K27ac indicates that epigenome rewiring occurs in p63 mutant keratinocytes.

To investigate the underlying mechanisms of enhancer re-distribution, a *de novo* motif scan was performed. We detected the p53/p63 motif family as the top enriched motif among control-specific enhancers (Fig 4E and Table EV6A), consistent with our previous results that the impaired p63 binding resulted in loss of active enhancers (Fig 3A). In contrast, motif analyses of mutant-specific enhancers captured motifs of bZIP, TEA, HMG and Runt family TFs (Table EV6B), among which some are p63 co-regulators (Kouwenhoven et al., 2015a). These data suggest that aberrant recruitment of TFs induces gain of enhancers.

One scenario of aberrant recruitment of TFs may result from the abnormal upregulation of TFs in mutant keratinocytes. To assess this possibility, we examined differential expression of TFs in control and mutant keratinocytes. Among 1581 examined TFs (Saeed, Quintin et al., 2014), 106 TFs were downregulated and 103 were upregulated in p63 mutant keratinocytes (Fig EV6). Most bHLH and basic leucine zipper domain (bZIP) family TFs were downregulated in p63 mutant keratinocytes. The large C2H2 zinc finger Krueppel factors and the Homeobox_dom family were upregulated in p63 mutant keratinocytes (Fig EV6). Interestingly, most downregulated TFs had p63 BSs nearby, and therefore they are potential p63 direct targets. In contrast, fewer upregulated TFs had p63 BSs, indicating that deregulation of many of these genes may be via indirect mechanisms (Fig EV6). To predict candidate TFs that are potentially bound to mutant-specific enhancers, we used two criteria: i) TFs whose binding motifs were enriched in mutant-specific enhancers (Fig 4E); ii) TFs that had upregulated gene expression in all three p63 mutant keratinocytes at the proliferation stage (day 0) shown by RNA-seq data (Fig EV6A). Among TFs that were consistently up-regulated in mutant keratinocytes, RUNX1and SOX4 belong to the TF families whose motifs were enriched in mutant-specific enhancers. We performed RT-qPCR validation to confirm the higher expression of SOX4 in p63 mutant keratinocytes (Fig EV6B). ChIP-qPCR of SOX4 also confirmed a number of SOX4 BSs with higher binding signals in R304W mutant keratinocytes compared to control keratinocytes (Fig EV6C). Interesting, one of the candidate TFs RUNX1 is a known p63 target (Masse, Barbollat-Boutrand et al., 2012) and a potential co-regulator that had many lost p63 BSs in the gene locus and was consistently upregulated in p63 mutant keratinocytes (Figs 3E and 5D, Fig EV6A).

**Figure 5.**
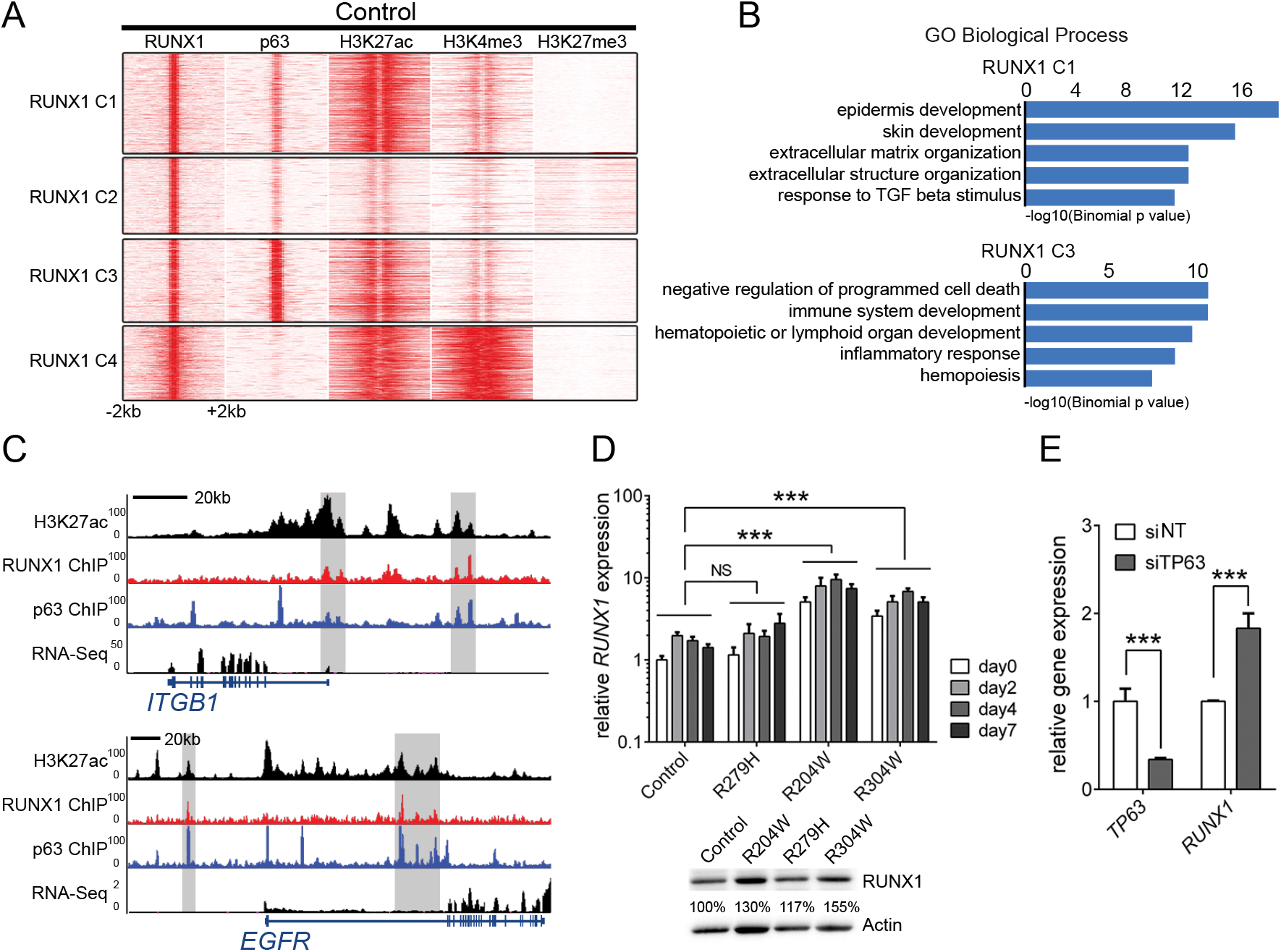
RUNX1 is a co-regulator and target gene of p63. A. K-means clustering of RUNX1 BSs detected in control keratinocytes. RUNX1 binding signals were shown in heatmaps in a 4-kb window with summits of RUNX1 BSs in the middle (*k* = 4, metric = Pearson). Color intensity represents normalized read counts. B. GREAT-based GO biological process annotation of RUNX1 BSs in RUNX1 C1 and C3. C. Representative example of RUNX1 and p63 co-regulated genes *ITGB1* and *EGFR*. D. Validation of *RUNX1* gene expression by RT-qPCR and western blotting. In RT-qPCR analysis, relative gene expression was normalized to the reference gene *hARP*. Data are shown as mean ± standard deviation, n=2, two-way ANOVA, NS *P* value > 0.05, *** *P* value <0.001. E. Gene expression analysis by RT-qPCR of *TP63* and *RUNX1* expression in *TP63* knockdown (siTP63) compared to non-targeting siRNA (siNT) in control keratinocytes. Gene expression was normalized to the reference gene *hARP*. Data are shown as mean ± standard deviation, n=2, two-way ANOVA, ***P value < 0.001. See also Fig EV6 and Table S7.

### Deregulated p63 and RUNX1 cooperation contributes to transcriptional rewiring in mutant p63 keratinocytes

To characterize p63 and RUNX1 co-regulation, we performed RUNX1 ChIP-seq in control keratinocytes. K-means clustering of RUNX1 BSs in combination with p63 binding and histone modification profiles showed that RUNX1 and p63 preferentially co-bound in active enhancer regions. RUNX1 bound more frequently to active promoters marked by H3K4me3 (RUNX1-C4) than p63 (RUNX1-C1 and RUNX1-C3) (Fig 5A and Table EV7A). Genes near the co-regulated enhancers (RUNX1-C1 and RUNX1-C3) were mainly involved in ‘epidermis development’ and ‘programmed cell death’, respectively (Fig 5B). For example, RUNX1 and p63 co-bound to both the promoter region and a proximal enhancer of *ITGB1*, and to the intronic enhancer region of *EGFR* (Fig 5C). The observed upregulation of RUNX1 expression (Fig 5D) and loss of p63 binding in the *RUNX1* gene body in p63 mutant keratinocytes (Fig 3E) indicated that deregulation of RUNX1 expression is probably due to loss of p63 control. To further confirm this, we performed siRNA knockdown of p63 in control keratinocytes. Similar to the upregulated *RUNX1* expression in p63 mutant keratinocytes, *RUNX1* expression was significantly increased in p63 knockdown keratinocytes (Fig 5E).

Given the possibility that upregulated *RUNX1* expression leads to its aberrant recruitment to mutant-specific enhancers in p63 mutant keratinocytes, we performed genome-wide comparison of RUNX1 binding in control keratinocytes and R304W mutant keratinocytes (Fig 6A and Tables EV7B-D). Among all RUNX1 BSs in both control and p63 mutant keratinocytes, there were 7,918 RUNX1 BSs with higher binding signals and 7,898 sites with lower binding signals in R304W mutant keratinocytes, as compared to RUNX1 binding in control keratinocytes (Tables EV7B-F). RUNX1 BSs with increased binding signals in R304W mutant keratinocytes were more often located in active enhancer regions accompanied by increased H3K27ac signals (Fig 6B and Table EV7E), while those with decreased RUNX1 binding signals were more often located in promoter regions (Figs EV7A and B, Table EV7F).

**Figure 6.**
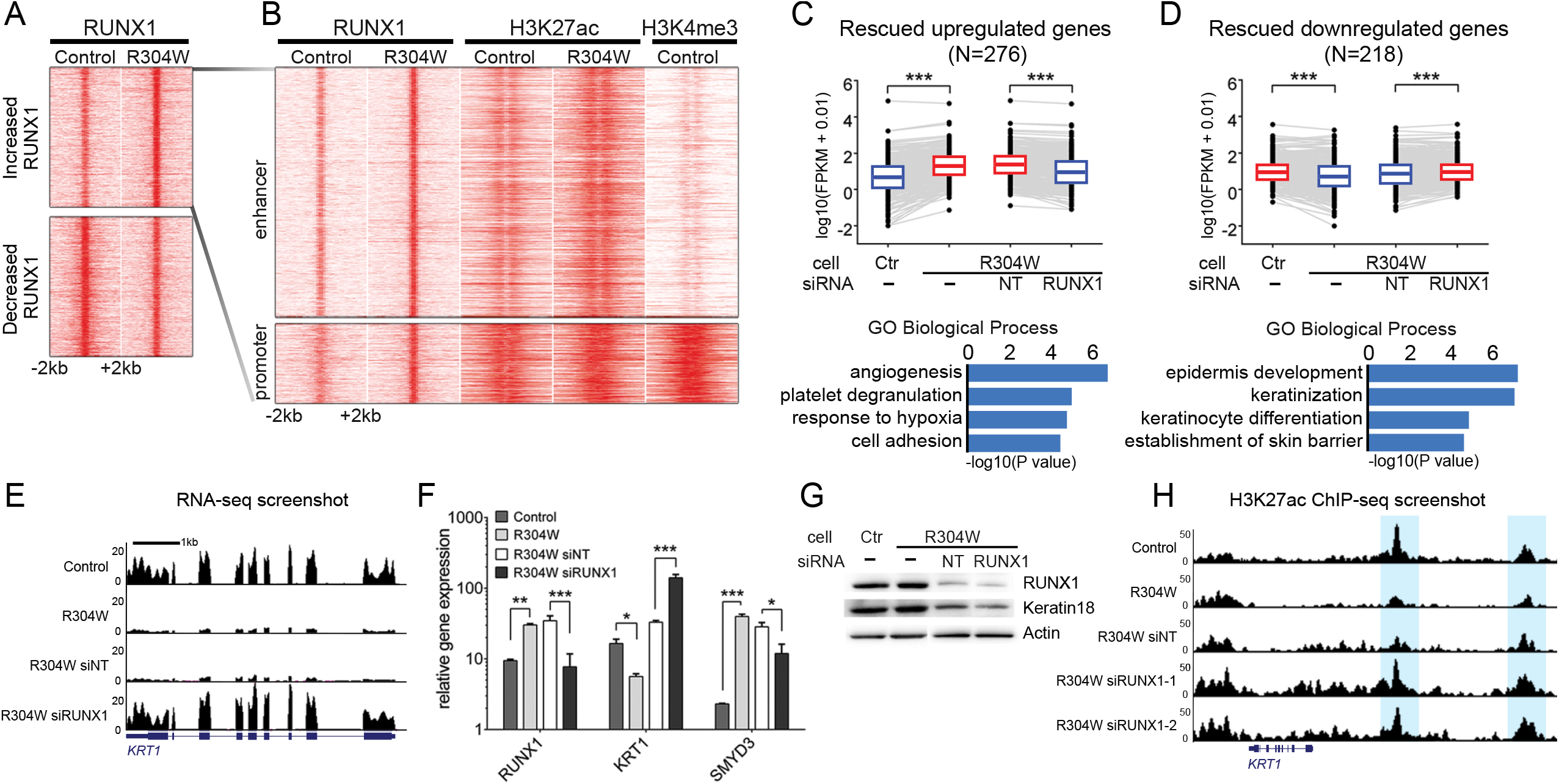
Increased RUNX1 binding contributes to gene deregulation. A. K-means clustering of RUNX1 BSs from control keratinocytes and R304W mutant keratinocytes were shown in heatmaps in a 4 kb window with summits of merged RUNX1 BSs in the middle (*k* = 2, metric = Pearson). Color intensity represents normalized read counts. B. Zoom-in re-clustering of increased RUNX1 BSs in R304W mutant keratinocytes, showing majority of RUNX1 BSs with increased signals are enhancers. C. Upregulated genes in R304W mutant keratinocytes that were rescued with knockdown of RUNX1 (siRUNX1) (*P* value <0.05). Expression (RNA-seq FPKM) of rescued genes under different conditions are shown as mean ± standard deviation. T-test, \*\*\**P* value < 0.001. DAVID-based GO biological process annotation is shown at the bottom. D. Downregulated genes in R304W mutant keratinocytes that were rescued with knockdown of RUNX1 (siRUNX1) (*P* value <0.05). Expression (RNA-seq FPKM) of rescued genes under different conditions are shown as mean ± standard deviation, T-test, \*\*\**P* value < 0.001. DAVID-based GO biological process annotation is shown at the bottom. E. UCSC genome browser screenshot of RNA-seq data at the gene loci of *KRT1* was shown, as examples of rescued genes by siRUNX1. F. Gene expression analyses by RT-qPCR of *RUNX1, KRT1 andSMYD3* upon RUNX1 knockdown in R304W mutant keratinocytes. Relative gene expression of these genes in control keratinocytes was normalized to the reference gene *hARP*. Data are shown as mean ± standard deviation, n=2, two-way ANOVA, \**P* value < 0.1, \*\**P* value < 0.01, \*\*\**P* value < 0.001. G. Western blotting analysis of RUNX1 and KRT18 were performed with protein extracts from control, R304W mutant keratinocytes, and R304W mutant keratinocytes treated with siNT or siRUNX1. Actin was used as the loading control. H. The UCSC genome browser screenshot of H3K27ac ChIP-Seq tracks at gene loci of *KRT1*, as examples of rescued enhancers, highlighted in blue, upon RUNX1 knockdown (siRUNX1) in R304W mutant keratinocytes. See also Fig EV7 and Table EV9.

To test whether increased RUNX1 expression is responsible for deregulating gene expression in p63 mutant keratinocytes, we performed RUNX1 knockdown in R304W mutant keratinocytes. As the proper RUNX1 expression level is important for cell proliferation (Hoi, Lee et al., 2010, Masse et al., 2012), we carefully titrated RUNX1 siRNA oligonucleotides in R304W mutant keratinocytes to achieve a similar level of RUNX1 expression to that in control keratinocytes. RNA-seq analyses showed that the overall gene expression pattern of R304W mutant keratinocytes treated with RUNX1 siRNA oligonucleotides (siRUNX1) was more similar to that of control keratinocytes in PCA, as compared to that of R304W mutant keratinocytes treated with the non-targeting siRNA (siNT) (Fig EV7C). Among the 3294 upregulated genes in R304W mutant keratinocytes compared to control keratinocytes, 276 genes were significantly rescued, having downregulated expression upon siRUNX1 treatment in R304W mutant keratinocytes (Fig 6C, Tables EV9A and B), e.g. *KRT8* (Fig EV7E). Many of the rescued genes were involved in adhesion and other non-epithelial functions such as ‘angiogenesis’, ‘platelet degranulation’. Importantly, among the 3187 downregulated genes in R304W mutant keratinocytes compared to control keratinocytes, 218 genes were significantly rescued, with upregulated expression upon siRUNX1 in R304W mutant keratinocytes (Fig 6D, Tables EV9C and D), for example, *KRT1* and *HES5* (Fig 6E and Fig EV7D). Many of these genes are important for epidermis development and keratinocyte differentiation. Partial but significant rescues by siRUNX1 was expected, as deregulated genes caused by loss of p63 binding in p63 mutant keratinocytes could not simply be rescued by RUNX1 knockdown. RT-qPCR and western blotting experiments validated rescued gene expression of *KRT1, SMYD3, and KRT18* in R304W mutant keratinocytes upon siRUNX1 (Figs 6F and G). To further investigate whether siRUNX1 can also partially rescue the enhancer landscape in R304W mutant keratinocytes, we performed H3K27ac ChIP-seq in these cells upon siRUNX1. Indeed, rescues at some enhancers were observed with a high reproducibility (Fig EV7F), e.g. the enhancers nearby *KRT1* (Fig 6H). Taken together, our data suggest that reversing upregulated RUNX1 expression can partially rescue deregulated gene expression and the enhancer landscape in R304W mutant keratinocytes.

## Discussion

The master regulator role of p63 in epidermal development has been established by many studies using *in vitro* and *in vivo* models (Browne et al., 2011, Mills et al., 1999, Yang et al., 1999). Recent studies also shed lights on p63 function in regulating the chromatin and enhancer landscape in epidermal keratinocytes (Bao et al., 2015, Cavazza et al., 2016, Kouwenhoven et al., 2015a, Rinaldi et al., 2016). However, it remains unclear how p63 mutations associated with developmental disorders affect the chromatin landscape and gene expression that contribute to the disease. In this study, we used EEC patient-derived skin keratinocytes carrying heterozygous p63 DNA-binding domain mutations as the cellular model to characterize the global gene regulatory alteration. We showed that the epidermal cell identity was compromised in p63 mutant keratinocytes, as indicated by downregulated epidermal genes and upregulated non-epithelial genes.

Consistent with the role of p63 in controlling the enhancer landscape during keratinocyte differentiation, deregulated gene expression was accompanied with an altered enhancer landscape. Loss of p63 binding led to reduced p63-bound active enhancers, while abnormally induced active enhancers were bound by deregulated p63 co-regulators such as RUNX1. Reducing expression levels of overexpressed RUNX1 in p63 mutant keratinocytes could partially rescue the deregulated gene expression as well as deregulated enhancers. Our data suggest an intriguing model that rewiring of the enhancer landscape, contributed by both loss of p63-bound active enhancers and gain of active enhancers induced by overexpressed p63 co-regulators such as RUNX1, gives rise to gene deregulation and phenotypes of EEC syndrome (Fig 7).

**Figure 7.**
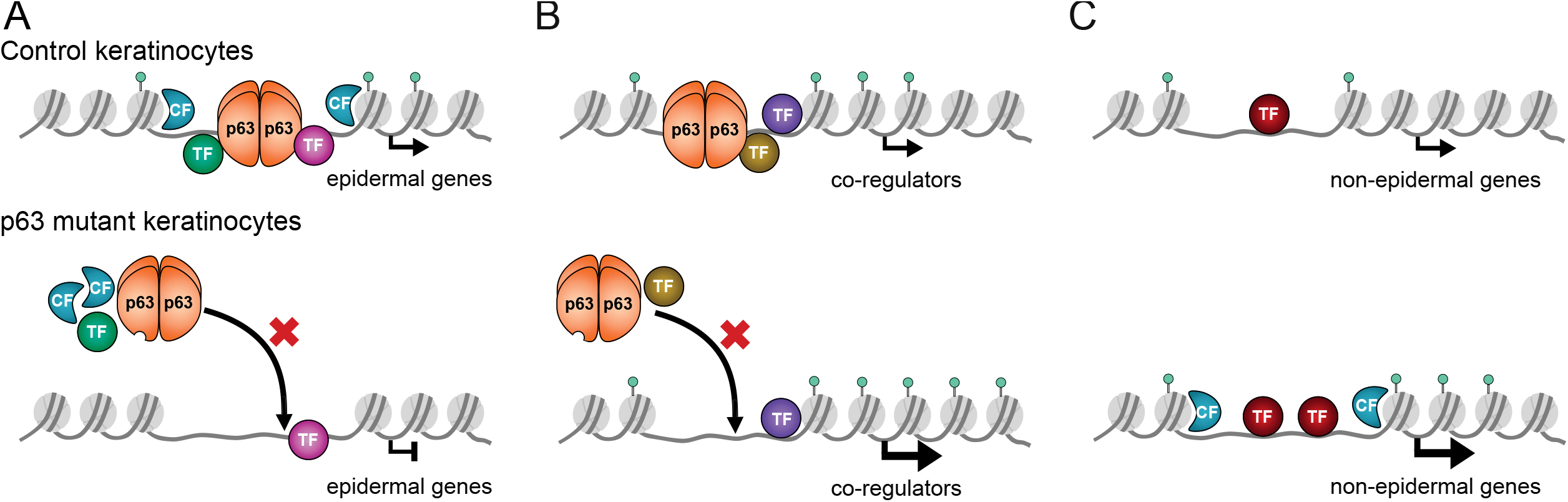
Model of p63-controlled gene regulation and cell identity in control and p63 mutant keratinocytes. A. p63-mediated activation of epidermal genes. In control keratinocytes, p63 activates epidermal genes by binding to nearby active enhancers, in cooperation with co-regulating TFs and chromatin factors (CFs). In p63 mutant keratinocytes, mutant p63 cannot bind to these enhancers, which results in downregulation of epidermal genes. B. p63-mediated fine tuning of its co-regulators. In control keratinocytes, p63 fine-tunes the expression of its co-regulators such as RUNX1 and SOX4, probably by recruiting repression complexes. In p63 mutant keratinocytes, mutant p63 cannot bind to these enhancers and lose the fine-tuning control. C. Indirect activation of enhancers due to lack of p63 control. Overexpressed p63 co-regulators such as RUNX1 can bind to less controlled open chromatin regions and enhancers to activate non-epidermal genes in p63 mutant keratinocytes.

Many epidermal genes were downregulated in mutant keratinocytes, consolidating the key role of p63 in epidermal development (Kouwenhoven et al., 2015b). In addition, upregulated mesenchymal genes and neuronal genes in mutant keratinocytes (Fig 1D) suggest that these cells have less defined epidermal cell fate and is consistent with previous findings (Barton et al., 2009, De Rosa, Antonini et al., 2009, Shalom-Feuerstein et al., 2011). For example, many mesodermal related genes, such as *ACTA2, ENC1, TPM2*, and *COL4A1*, which were upregulated during epidermal commitment of p63-depeleted embryonic stem cells (Shalom-Feuerstein et al., 2011) were also upregulated in p63 mutant keratinocytes (Table EV1E). These findings are also in agreement with the gene co-expression network analyses that showed two modules ‘keratinization’ and ‘nucleosome assembly’ associated with downregulated genes in mutant keratinocytes were connected with the scattered upregulated genes involved in ‘extracellular matrix organization’ (Fig 1E). This suggests that gene expression in p63 mutant keratinocytes deviates from the proper epidermal cell fate to establish a new differentiation direction through chromatin remodeling processes. In addition to directly regulating chromatin remodeling factors (Fig EV2B), p63 has been shown to be important for the chromatin landscape by establishing the enhancers (Bao et al., 2015, Kouwenhoven et al., 2015a). Consistently, we showed in this study that p63 motif was most significantly enriched in active enhancers at early differentiation stages (Fig 2E), supporting the pivotal role of p63 in regulating enhancers to activate epidermal differentiation genes, especially at the initiation stage (Bao et al., 2015, Kouwenhoven et al., 2015a). Interestingly, we observed persistent higher p63 expressions till the end of differentiation in p63 mutant keratinocytes, as compared to the control keratinocytes. The high expression of p63 at the end of differentiation is consistent with differentiation defects observed in p63 mutant keratinocytes.

In addition to its activator role for epidermal genes, it has been shown that p63 can function as a repressor to repress p21 (*CDKN1A*) through recruiting HDAC1/2 (LeBoeuf et al., 2010, Ramsey, He et al., 2011). RUNX1, a co-regulator of p63, is regulated by p63 in a complex fashion. Depending on the differentiation state of keratinocytes, p63 activates or represses *RUNX1* expression (Masse et al., 2012). In our analyses, we detected upregulation of *RUNX1* in proliferating mutant keratinocytes (Fig 3E and Fig 5D), consistent with the repressor role of p63 for these genes. Interestingly, instead of repressive histone modification marks such as H3K27me3 and H3K9me3, we observed that the active enhancer mark H3K27ac co-localized with p63 BSs at the RUNX1 locus, suggesting that p63 acts as an activator, and that these genes are not at a completely repressed state. Furthermore, no H3K27me3 or H3K9me3 repression mark was found at any p63 BS (Kouwenhoven et al., 2015a), indicating that p63-bound enhancers are not repressed by the polycomb- or heterochromatin-related mechanisms. To reconcile the apparent contradiction that p63 binds to active enhancers to repress gene expression, we propose that p63 binds to these active enhancers to fine tune expression of these genes, maybe by recruiting repressors such as HDACs (LeBoeuf et al., 2010, Ramsey et al., 2011). When p63 expression or function is compromised, such as in p63 mutant keratinocytes or in p63 knockdown cells (Figs 5D and E), the fine tuning or repression mechanisms are relieved and the expression of these genes is enhanced. In this study, we showed that p63 binding loss and loss of active enhancers occurred at a genome-wide scale in patient keratinocytes carrying heterozygous EEC mutations (Fig 3A). This indicates that the mutant p63 protein has a dominant effect on DNA binding over the wild type p63 that is also present in the cell, because the p63 protein is DNA-binding competent as a tetramer (Dotsch, Bernassola et al., 2010, Serber, Lai et al., 2002). These findings corroborate the dominant negative model of EEC p63 mutations that has been proposed by several previous studies using *in vitro* approaches (Browne et al., 2011, Celli et al., 1999) and allele-specific knockdown of p63 mutant (Novelli, Lena et al., 2016).

Unexpectedly, we also observed a large number of gained active enhancers in all three p63 mutant keratinocytes (Figs 4A and B, Fig EV5). These mutant-specific enhancers were enriched for motifs of TFs that normally cooperate with p63 in keratinocytes (Fig 4E). Many of these TFs were deregulated in p63 mutant keratinocytes (Fig EV6). Some of them such as RUNX1 are direct p63 targets and upregulated in p63 mutant keratinocytes (Figs 3E and 5D).This indicates that rewiring of the transcriptional program is not only contributed by p63 binding loss and thereby loss of active enhancers but also by an indirect effect of altered expression of p63 co-regulating TFs. ChIP-seq analyses revealed an altered RUNX1 binding profile in p63 mutant keratinocytes (Fig 6A). The increased RUNX1 binding was associated with increased H3K27ac signals (Fig 6B). In general, the re-distribution of active enhancers correlated with changed gene expression in p63 mutant keratinocytes (Fig 4E). Intriguingly, reversing *RUNX1* expression upon siRUNX1 in mutant keratinocytes could partially rescue gene deregulation and the enhancer landscape (Figs 6C-H), indicating that overexpression of *RUNX1* in mutant keratinocytes contributes to the gene deregulation and is at least partially responsible for the differentiation defects. Taken together, at the molecular level, mutations in the p63 DNA-binding domain can give rise to an indirect gain-of-function effect via inducing aberrant binding of deregulated TFs to genome-wide enhancers (Fig 6A).

This gain-of-function model of p63 mutations bears an interesting resemblance to that of p53 mutations that give rise to cancers (Zhou, Wang et al., 2014, Zhu, Sammons et al., 2015). Belonging to the same gene family, p53 and p63 share high sequence and structural similarity, especially in their DNA-binding domain (Celli et al., 1999, Courtois, de Fromentel et al., 2004). The amino acid residues in p63 such as R204, R279 and R304 that are mutated in EEC syndrome are also hot spots for mutations of p53 in cancer (Celli et al., 1999). It has been shown that some of these p53 mutants cannot bind to *bona fide* p53 targets, but instead, they cooperate with normal p53 co-regulators and bind to ectopic genomic sites to activate abnormal gene expression (Zhou et al., 2014, Zhu et al., 2015). Therefore, these p53 mutations have been proposed to act in both dominant negative and gain-of-function fashions, which is similar to the notion that we propose here for p63 EEC mutations. However, the most apparent difference between p53 and p63 mutations is that mutant p63 does not seemingly bind to ectopic genomic loci, as we did not observe any novel p63 BS in mutant keratinocytes (Fig 3A). The indirect gain-of-function action of p63 mutations is due to the overexpression of p63 co-regulating TFs such as RUNX1 and their induced aberrant recruitment to genomic loci, which leads to the abnormal gene expression. It should be noted that RUNX1-induced enhancers in p63 mutant keratinocytes were not *de novo* enhancers but showed quantitative differences in binding signals and enhancer activities. Similar to our experiments where we did not expect a full rescue upon RUNX1 knockdown in p63 mutant keratinocytes, as p63 binding loss could not simply be rescued by RUNX1 downregulation. We also do not expect that overexpression of RUNX1 would fully mimic p63 mutant keratinocyte phenotypes. However, it would be interesting to test whether overexpression of *RUNX1* in control keratinocytes can induce any differentiation defects. Furthermore, it is also of interest to further test the effect of other overexpressed p63 co-regulators, such as TFs of the bZIP, TEA and HMG families (SOX4) (Fig 4F) in the EEC disease mechanism. It is conceivable that knockdown of multiple such overexpressed TFs may rescue differentiation defects of p63 mutant keratinocytes to a better extent.

In conclusion, we identified a rewired enhancer landscape as a common mechanism in p63 mutant EEC keratinocytes. The enhancer rewiring includes loss of p63-bound active enhancers that regulate epidermal genes (Fig 7A) and gain of enhancers bound by overexpressed TFs that are normally fine-tuned by p63 (Figs 7B and C). This model is consistent with the proposed bookmarking role of p63 (Kouwenhoven et al., 2015a, Kouwenhoven et al., 2015b) in epithelial cells where p63 co-regulators play an important role in gene regulation. It is conceivable that, in p63 mutant EEC keratinocytes, the chromatin environment is less tightly controlled, and less-controlled TFs can therefore bind to more exposed enhancers. The rewired enhancer landscape gives rise to gene deregulation that contributes to the less-defined epidermal cell fate and skin phenotypes of the disease. It is known that the proper p63 function is essential in embryonic epithelial tissues such as the apical ectodermal ridge and the palatal epithelium for limb and palate development (Kouwenhoven et al., 2010, Thomason, Zhou et al., 2010). Our findings suggest that EEC mutations affect the epithelial cell fate of the embryonic epithelium during morphogenesis and development of related organs, and thereby contribute to limb malformation and cleft lip/palate. Taken together, the enhancer landscape rewiring contributes to the disease mechanism of p63 mutations, and may be common to many other diseases.

## Materials and Methods

### Ethics statement

All procedures for establishing and maintaining human primary keratinocytes were approved by the ethical committee of the Radboud university medical center (“Commissie Mensgebonden Onderzoek Arnhem-Nijmegen”). Informed consent was obtained from all donors of a skin biopsy.

### Human primary keratinocyte culture

Primary keratinocytes were established previously from skin biopsies of three EEC syndrome patients carrying heterozygous mutations in the p63 DNA-binding domain, R204W(van Bokhoven, Hamel et al., 2001), R279H(Brunner, 2002), and R304W(Celli et al., 1999), as well as of two healthy volunteers (Dombi23 and PCK19, referred to as Control) (Rheinwald & Green, 1977). As previously described(Rinne, Clements et al., 2008), primary keratinocytes were cultured in Keratinocyte Basal Medium (KBM, Lonza #CC-4131) supplemented with 100 U/mL Penicillin/Streptomycin (Gibco Life Technology #15140122), 0.1 mM ethanolamine (Sigma Aldrich #141–43–5), 0.1 mM O-phosphoethanolamine (Sigma Aldrich #1071–23–4), 0.4% (vol/vol) bovine pituitary extract, 0.5 μg/mL hydrocortisone, 5 μg/mL insulin and 10 ng/mL epidermal growth factor (Lonza #CC-4131). Medium was refreshed every other day. When cells were more than 90% confluent (day 0), differentiation was induced by depletion of growth factors in addition to cell contact inhibition, as described previously (Van Ruissen, De Jongh et al., 1996). Cells were collected at four differentiation stages, proliferation (day 0), early differentiation (day 2), mid differentiation (day 4), and late differentiation (day 7) for subsequent experiment. No mycoplasma contamination is found during cell culture.

### RNA extraction and quantitative real-time reverse transcription PCR (RT–qPCR)

Total RNA was isolated using the NucleoSpin RNA kit (MACHEREY-NAGEL #740955.250) and quantified with NanoDrop. cDNA synthesis from 1 μg freshly prepared total RNA was carried out using the iScript cDNA synthesis kit (Bio-Rad #170-8891). Reverse transcription quantitative PCR (RT-qPCR) primers were designed using Primer3 (http://frodo.wi.mit.edu), to obtain exon spanning primers wherever possible. Each primer set has been tested for its linear amplification dynamic range. RT-qPCRs were performed in the CFX96 Real-Time system (Bio-Rad) by using iQ SYBR^®^ Green Supermix (Bio-Rad #1708887) according to the manufacturer’s protocol. The human acidic ribosomal protein (*hARP*) or glucuronidase beta (*GusB*) was used as the housekeeping gene for normalization. Differences in the expression of each gene during differentiation (relative expression) were calculated by 2ΔΔCt method (Livak & Schmittgen, 2001). Sequences of all RT-qPCR primers (Biolegio BV, Nijmegen, the Netherlands) were provided in Table EV1C.

### Western blotting

About 20,000 keratinocytes under each condition were lysed in 50μl RIPA lysis buffer. After 30s sonication with Bioruptor^®^ Pico, the lysates were rotated in lysis buffer for 30min before a centrifuge at the maximum speed (17000rpm) for 30 minutes at 4°C. The supernatant was collected as protein extracts for western blotting. Total protein concentration of the supernatant was determined with BCA assay, and a total of 12.5 μg protein was loaded for each sample. The actin antibody (Sigma-Aldrich #clone AC-15, 1:100,000) was used to control equal protein loading. Protein extracts were run on SDS PAGE and transferred to PVDF membranes using the NuPAGE system (Life Technologies). LumiGLO (Cell Signaling Technology Inc.) was used for chemiluminescent detection by the Bio-Rad Universal Hood Gel Imager (Bio-Rad Laboratories). Antibodies used in this study include LOR (Babco COVANCE #PRB-145P, 1:2500), RUNX1 (Abcam #ab23980; 1:50), K18 (Merck #MAB3234, 1:500). All the original blots were provided in Table EV11.

### RNA-seq and analysis pipeline

RNA-seq experiment was performed as described previously (Kouwenhoven et al., 2015a) with the starting material of 500 ng total RNA, to obtain double-strand cDNA (ds-cDNA). After purification with the MinElute Reaction Cleanup Kit (Qiagen #28206), 3 ng ds-cDNA was processed for library construction using KAPA Hyper Prep Kit (Kapa Biosystems #KK8504) according to the standard protocol except that a 15-minute USER enzyme (BioLab # M5505L) incubation step was added before library amplification. The prepared libraries were quantified with the KAPA Library Quantification Kit (Kapa Biosystems #KK4844), and then sequenced in a paired-ended manner using the NextSeq 500 (Illumina) according to standard Illumina protocols.

Sequencing reads were aligned to human genome assembly hg19 (NCBI version 37) using STAR 2.5.0 (Dobin, Davis et al., 2013) with default options. Briefly, STAR has the option to generate in-house reference genome from the genome fastq file. In this study, hg19 genome was used to generate the in-house reference genome with the following command: STAR --runThreadN 8 --runMode genomeGenerate --genomeDir directory/ --genomeFastaFiles hg19.fa --sjdbGTFfile

Homo_sapiens.GRCh37.75.gtf --sjdbOverhang 100. Then STAR was run and it automatically generated read-counts directly. A detailed summary of RNA-seq data generated in this study was shown in Table EV10A. For data visualization, wigToBigWig from the UCSC genome browser tools was used to generate bigwig files and uploaded to UCSC genome browser. Genes with the mean of DESeq2-normalized counts (“baseMean”)> 10 were considered to be expressed. Differential gene expression (adjusted P value < 0.05) and principal-component analysis were performed with the R package DESeq2 using read counts per gene (Love, Huber et al., 2014). Hierarchical clustering was performed based on log10 (FPKM+0.01). Functional annotation of genes was performed with DAVID (Huang da, Sherman et al., 2009). For Weighted Gene Co-expression Network Analysis (WGCNA (Langfelder & Horvath, 2008)), only high variance genes (adjusted *P* value < 0.01, sum of baseMean > 100, 3162 genes) between control keratinocytes and p63 mutant keratinocytes during differentiation were included. WGCNA clustering within p63 mutant keratinocyte samples was performed using power of 26 and the minimum module size of 15. A total of 16 co-expression modules were identified based on gene co-expression patterns (Table EV1J). To visualize the gene network organization, only nodes (genes) with connectivity weight > 0.1 were kept (Table EV1K). Cytoscape (Smoot, Ono et al., 2011) was used for gene network visualization.

### ChIP-seq and analysis pipeline

Chromatin for ChIP was prepared as previously described(Kouwenhoven et al., 2010). ChIP assays were performed following a standard protocol (Novakovic, Habibi et al., 2016) with minor modifications. On average, 0.5M keratinocytes were used in each ChIP. For histone marks, 2x ChIP reactions were pooled to prepare 1x ChIP-seq sample; for transcription factors, 4x ChIP reactions are pooled to prepare 1 ChIP-seq sample. Antibodies against H3K27ac (Diagenode #C15410174, 1.2 μg), H3K4me3 (Diagenode #C15410003, 1 μg), H3K27me3 (Diagenode #C15410069, 1.5 μg), p63 (Santa Cruz #H129, 1 μg, recognizing the C-terminal α tail of p63) and RUNX1 (Abcam #ab23980, 4 μg) were used in each ChIP assay. Resulted DNA fragments from four independent ChIP assays were purified and subjected to a ChIP-qPCR quality check. Primers used for ChIP-qPCR were listed in Table EV1C. Afterwards 5ng DNA fragments were pooled and proceeded on with library construction using KAPA Hyper Prep Kit (Kapa Biosystems #KK8504) according to the standard protocol. The prepared libraries were then sequenced using the NextSeq 500 (Illumina) according to standard Illumina protocols.

Sequencing reads were aligned to human genome assembly hg19 (NCBI version 37) using BWA (Li & Durbin, 2009). Mapped reads were filtered for quality, and duplicates were removed for further analysis. A detailed summary of ChIP-seq data generated in this study was shown in Table EV10B. In addition. The bamCoverage script was used to generate and normalize bigwig files with the RPKM formula. The peak calling was performed with the MACS2 (Zhang, Liu et al., 2008) against a reference input sample from the same cell line with standard settings and a q value of 0.05. The broad setting (--BROAD) was enabled for detecting H3K27me3 peaks. Only peaks with a *P* value < 10e-5 were used for differential analysis with MAnorm (Shao et al., 2012). The log2 ratio of read density between control keratinocytes and each mutant keratinocytes (M value) was plotted against the average log2 read density (A value) for all peaks. By merging all the significantly higher H3K27ac peaks in control keratinocytes in each comparison, there are in total 15,057 peaks which are named as ‘control-specific enhancers’; by merging all the significantly higher H3K27ac peaks in each p63 mutant keratinocytes, there are in total 17,931 peaks which are named as ‘mutant-specific enhancers’. Association of peaks to genes and associated GO annotation were performed with GREAT (McLean et al., 2010), with the ‘single nearest gene within 1 Mb’ association rule. *P* values were computed with a hypergeometric distribution with FDR correction. k-means clustering and heat map and band plot generation were carried out with a Python package fluff (Georgiou & van Heeringen, 2016). H3K27ac ChIP-seq analyses of control and mutant keratinocytes, including those performed in siRUNX1 experiments, were performed in duplicates. The ‘plotCorrelation’ function from the deepTools package was used and the Pearson correlation coefficient was calculated accordingly (Ramírez, Dündar et al., 2014).

### ChomHMM analysis

Chromatin states were characterized using ChromHMM v1.11 (Ernst & Kellis, 2012). The peak files from three tracks (H3K27me3, H3K4me3, and H3K27ac) across four stages were used as input. The six-emission state model was determined to be optimal which could sufficiently capture the biological variation in co-occurrence of chromatin marks. The segmentation files of the six emission states per stage were binned into 200 bp intervals. An M × N matrix was created, where M corresponds to the 200 base pair intervals and N to the developmental stages (N=4). Each element x (m, n) represents the chromatin state of interval m at stage n. For each chromatin group, occurrences were counted per stage N (Table EV3B). The changes between stage N and N+1 were plotted pair-wise using Sankey diagrams (http://sankeymatic.com/).

### Motif analysis

Previously described p63scan algorithm was used for p63 motif evaluation(Kouwenhoven et al., 2010). HOMER (http://homer.salk.edu/homer/motif/) was used for motif scan against corresponding background sequences. For example, enriched motifs in regions migrating from unmodified regions to active enhancers were screened against regions remaining unmodified; enriched motifs in regions migrating from active enhancers to unmodified regions were screened against regions remaining active enhancers; and enriched motifs in control-specific enhancers were screened against mutant-specific enhancers.

### siRNA nucleofection

Nucleofection was performed as described previously (Mulder, Wang et al., 2012) using the Amaxa 96- well shuttle system (Lonza, program FF113). In short, keratinocytes were harvested with Accutase^®^ solution and resuspended in nucleofector buffer SF (Lonza). Each 20 μL transfection reaction contained 200,000 cells mixed with 2 μM validated siRNA (Silencer Select siRNAs, Ambion/Applied Biosystems). siRNA information was listed in Table EV8. In siRUNX1 experiments, siRUNX1-1 oligo was used for the RNA-seq experiment and siRUNX1-2 oligo was used for the western blot experiment. Both siRUNX1 oligoes were used in the RT-qPCR and H3K27ac ChIP-seq experiments. After transfection, the samples were incubated for 10 minutes at room temperature before resuspension in KBM and seeding at 50.000 cells per well (12 well plate). Medium was refreshed each day for the indicated periods till the end of experiments.

### Statistics and reproducibility

Data are expressed as mean ± standard deviation error of the mean unless otherwise specified. Data set statistics were analyzed using the GraphPad Prism software. Differences under *P* < 0.05 were considered statistically significant, NS *P* value > 0.05, * *P* value <0.05, ** *P* value <0.01, *** *P* value <0.001. Gene expression analysis by RT-qPCR was performed in biological duplicates (n>=2); data are shown as mean ± standard deviation, two-way ANOVA. The comparison of gene expression (fold change) was analyzed with the unpaired T-test. Hypergeometric test was performed with the online tool GeneProf. The comparison of gene expression after siRNA knockdown was performed with T-test. Other statistical methods used in this study were specified in the Figure legends.

### Data availability

To review our complete dataset GEO accession GSE98483 (https://www.ncbi.nlm.nih.gov/geo/query/acc.cgi?acc=GSE98483). To review our complete dataset in Genome Browser, please go to https://genome.ucsc.edu/cgi-bin/hgTracks?hgS_doOtherUser_submit&hgS_otherUserName_Jieqiong_20QU&hgS_otherUserSessionName_hg19_p63_RUNX1_Jieqiong

Genomic data of normal control individuals have been deposit in the GEO database (GSE98483), and those of p63 patients will be deposit in dbGaP database with controlled access. All data supporting the findings of the study and in-house codes are available on request.

## Acknowledgements

We thank Gert Jan Veenstra for discussions and suggestions on the project. We thank Georgios Georgiou, Thomas te Wierik, and Guoqiang Yi for their help with data analyses, Hanna Niehues and Gijs. A.C. Franken for providing technical support of experiments, Eva Janssen-Megens, Siebe van Genesen and Rita Bylsma for operating the Illumina analyzer and initial data output. We thank the ENCODE Consortium for sharing their data.

## Author contributions

JQ and HZ conceived and designed the experiments. JQ, ST, JPHS, ENK, MO, CL, HGS, HvB, KM, HZ wrote and revised the manuscript. JQ, ST, JPHS, ENK and EHvB performed the experiments. JQ, MO and HZ analyzed the data.

## Expanded View Figures

**Figure EV1.**
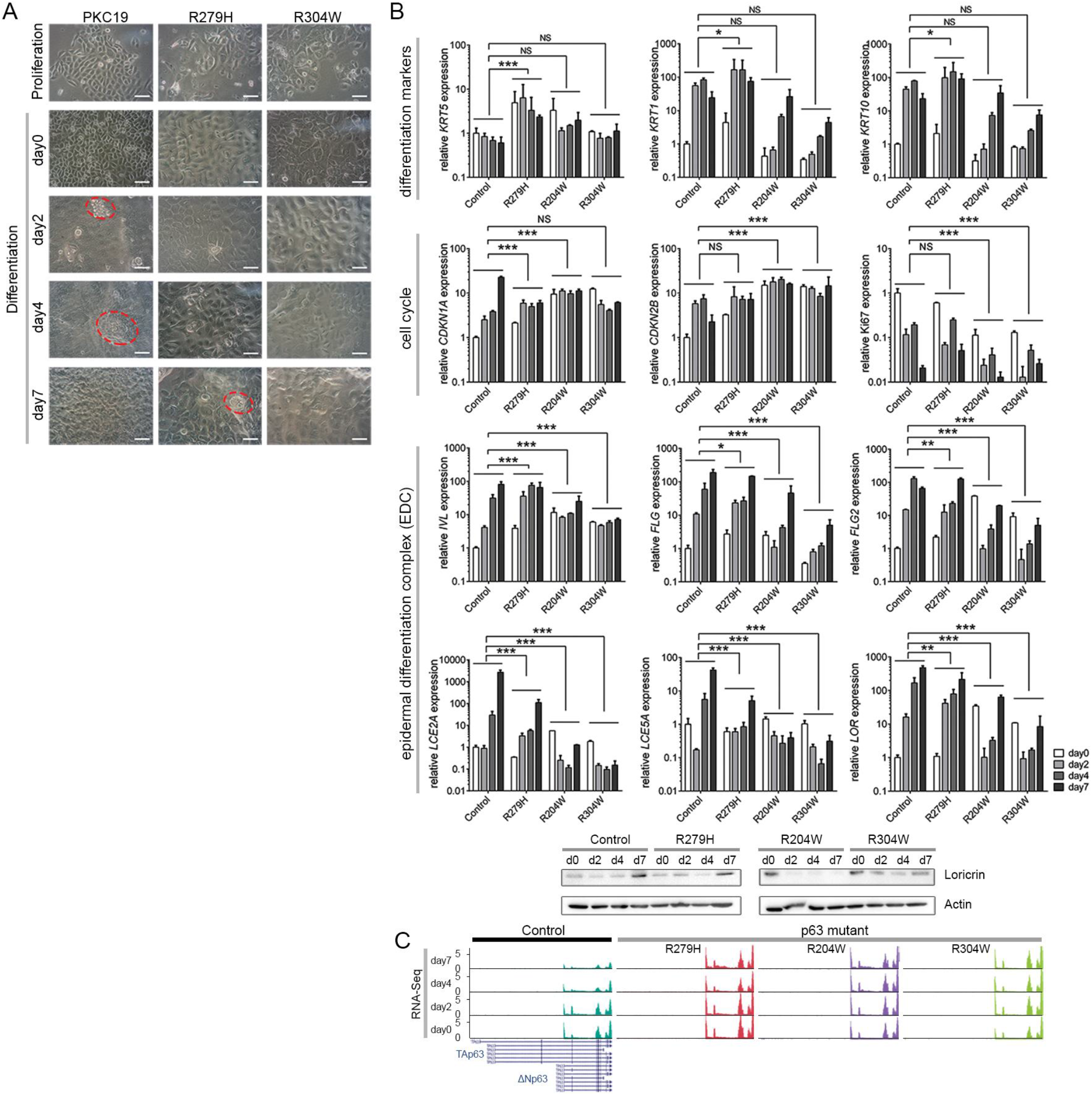
Differentiation defects in p63 mutant keratinocytes, related to Figure 1. A. Morphology difference observed between control and p63 mutant keratinocytes. Multi-layer structures are indicated with red circles. All scale bars, 50 μm. B. Gene expression validation analysis by RT-qPCR and western blot. Several differentiation markers, cell cycle and epidermal differentiation complex (EDC) genes, were validated by RT-qPCR at the mRNA level. Gene expression was normalized to the expression of *GusB* (reference gene). Primer information is provided in Table EV1C. Data are shown as mean ± standard deviation, *n*=2, two-way ANOVA, NS *P* value > 0.05, * *P* value <0.05, ** *P* value <0.01, *** *P* value <0.001. Expression of the epidermal marker gene, Loricrin, was also validated by western blot at the protein level. Actin was used as a loading control. C. A UCSC genome browser screenshot of RNA-Seq tracks at the *TP63* gene loci. Expression of TA-specific exons were not detected.

**Figure EV2.**
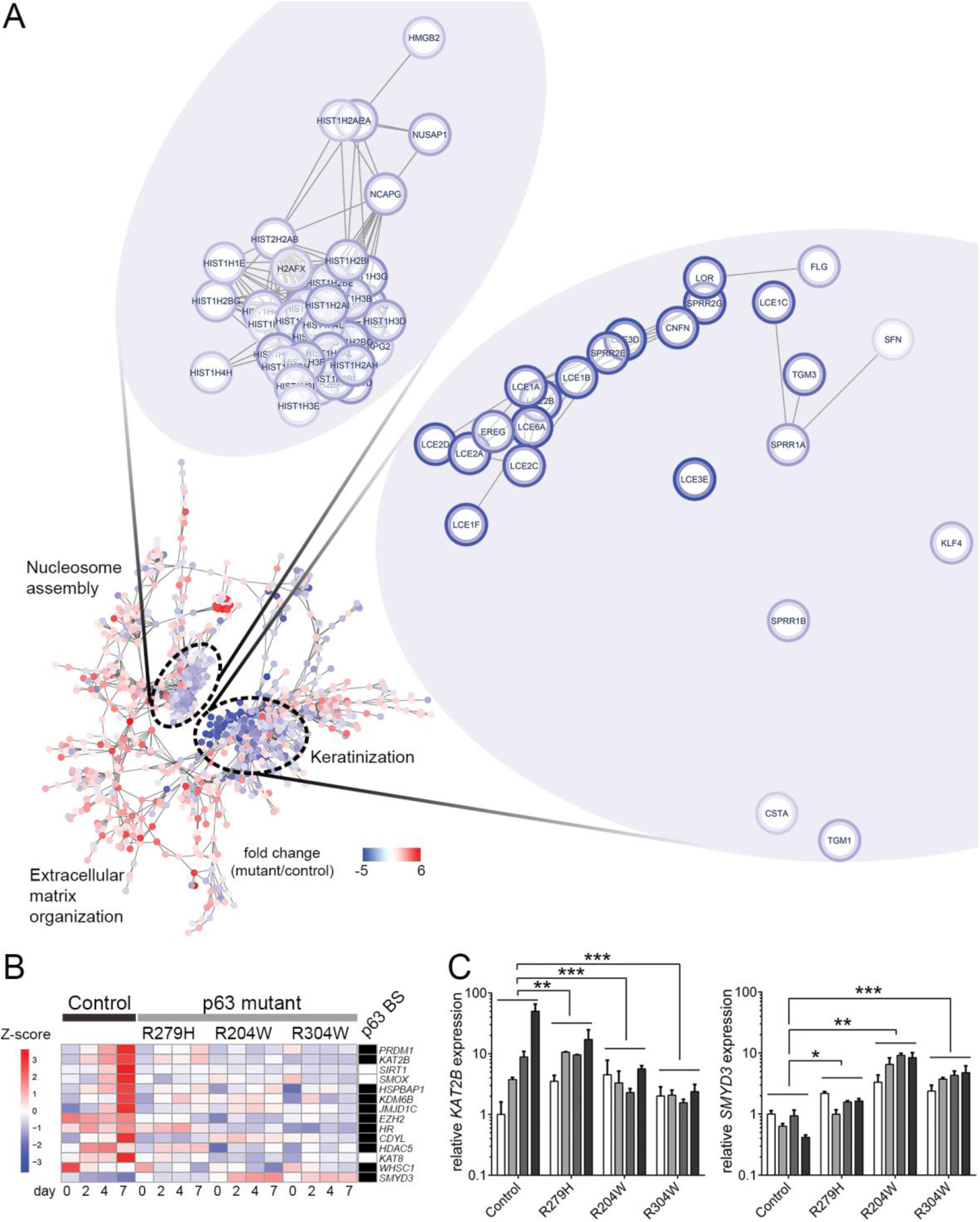
Deregulated gene network and chromatin regulators in p63 mutant keratinocytes, related to Figure 1. A. Visualization of representative hub genes in ‘nucleosome assembly’ module and ‘keratinization’ module using the Cytoscape Network Analyzer. B. Gene expression heatmap of differentially regulated chromatin factors (*P* value <0.01 and fold change >2) in p63 mutant keratinocytes. Nearby p63 binding sites (p63 BSs) of these genes are indicated, black, with at least one p63BS; white, without p63BS. Z-score is calculated based on log10 (FPKM+0.01) of each gene. C. Gene expression analysis by RT-qPCR of *KAT2B* and *SMYD3*. Relative gene expression is shown based on normalization to expression the reference gene *GusB*. Data are shown as mean ± standard deviation, *n*=2, two-way ANOVA, * *P* value <0.05, ** *P* value <0.01, *** *P* value <0.001.

**Figure EV3.**
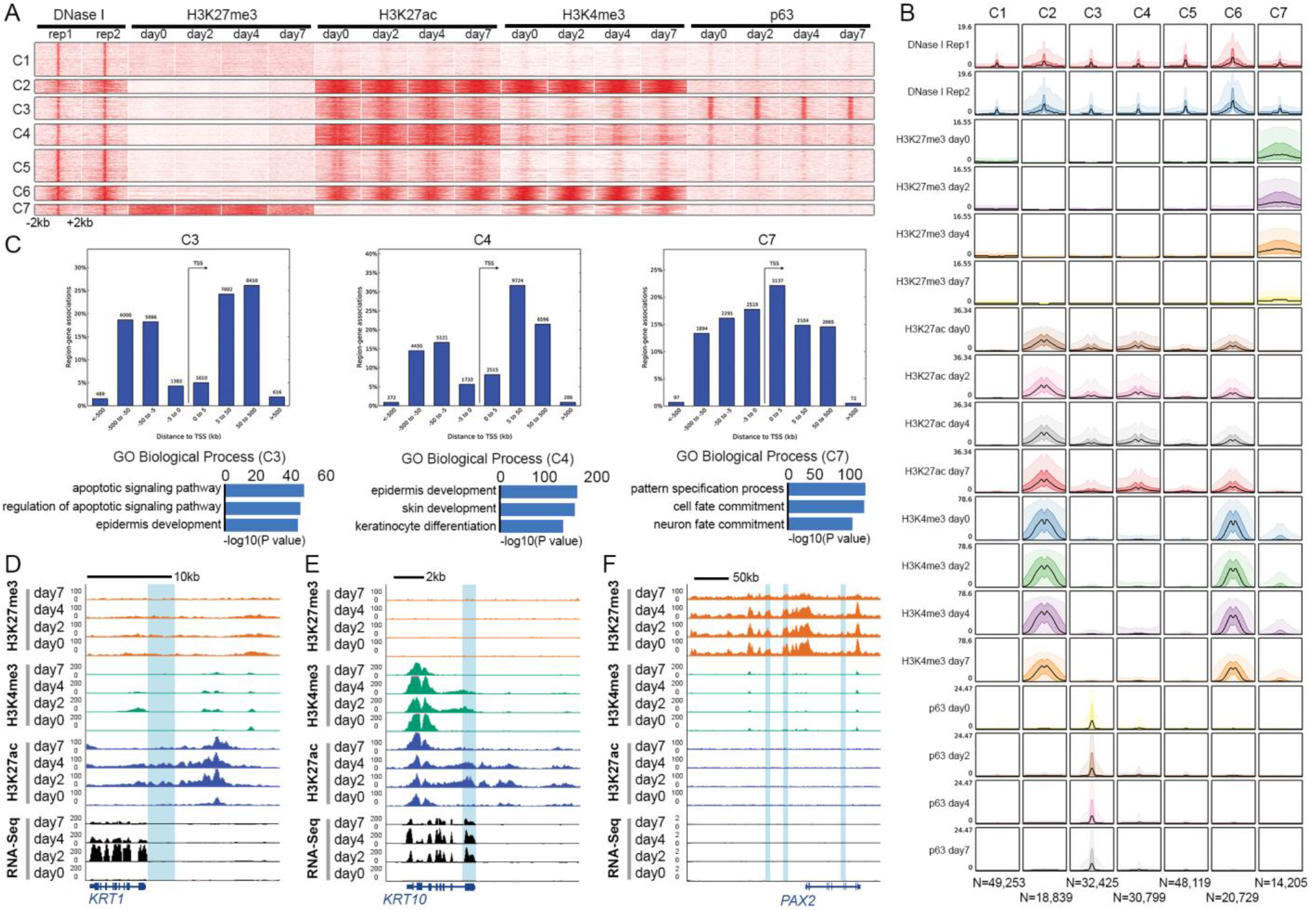
Characterization of the epigenetic landscape during keratinocytes differentiation, related to Figure 2. A. Signal heatmap of several histone marks during keratinocyte differentiation at open chromatin regions detected by DNase I Hypersensitivity Sites (DNase I). K-means clustering groups DNase I peaks into seven classes (*k* = 7, metric = Pearson). Heatmaps are plotted in a 4 kb window with summits of DNase I peaks in the middle. Color intensity represents normalized read counts. B. Band plots that quantifies the signals in the heatmap shown in A. The number of regions in each cluster is illustrated at the bottom. C. Characterization of the active enhancer clusters and heterochromatin cluster. Top bar charts, the distance between detected regions and their putatively regulated genes; bottom bar charts, top three enriched GO biological processes annotated with GREAT. D. An example of chromatin state dynamics (unmodified regions and active enhancers) during differentiation at the gene loci of *KRT1*. E. An example of chromatin state dynamics (unmodified regions and active promoters) during differentiation at the gene loci of *KRT10*. F. An example of chromatin state dynamics (heterochromatin regions and unmodified regions) during differentiation at the gene loci of *PAX2*.

**Figure EV4.**
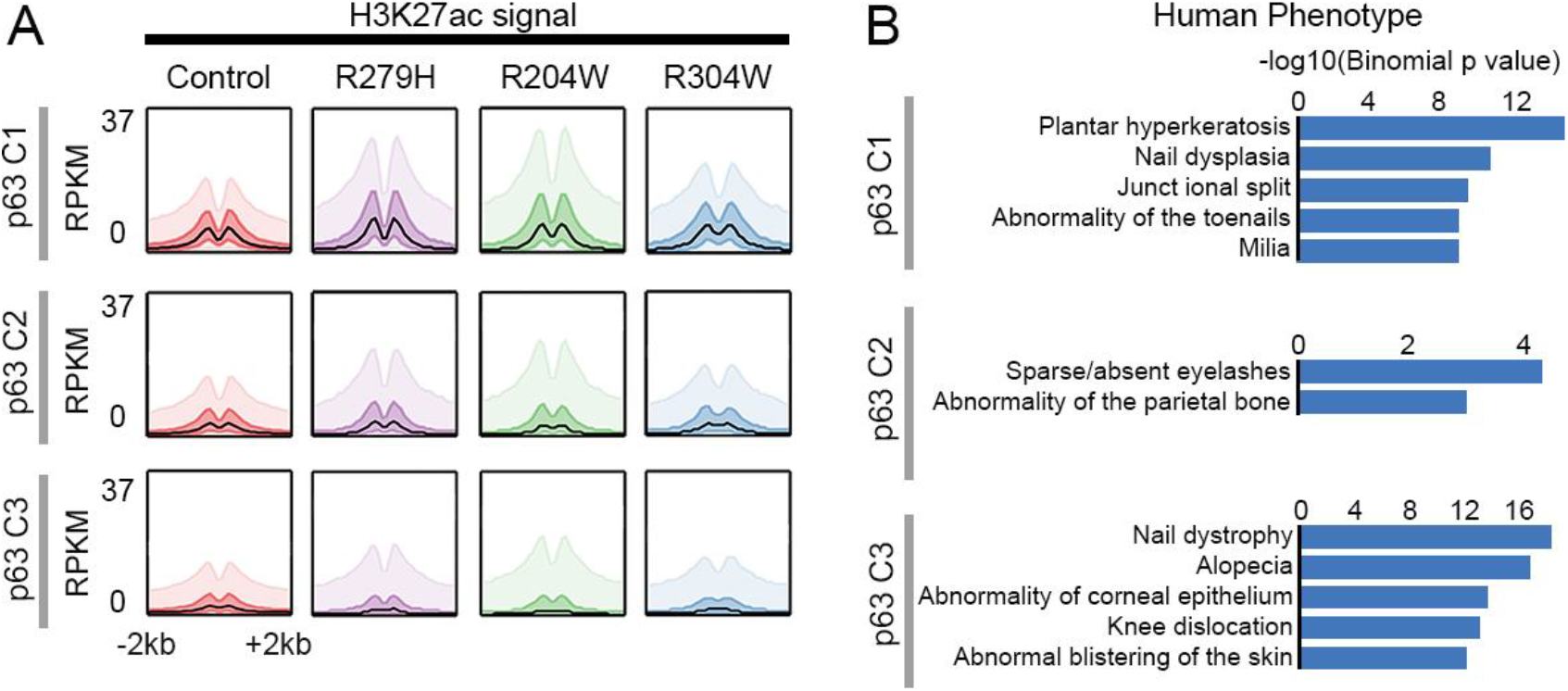
H3K27ac signal and human phenotype related to p63 BSs in three different clusters, related to Figure 3. A. Band plots of H3K27ac ChIP-seq signals in each p63 cluster. The median enrichment was visualized as a black line while 50% and 90% ranges were depicted in lighter color, respectively. B. GREAT-based human phenotype annotation of p63 BSs in each cluster.

**Figure EV5.**
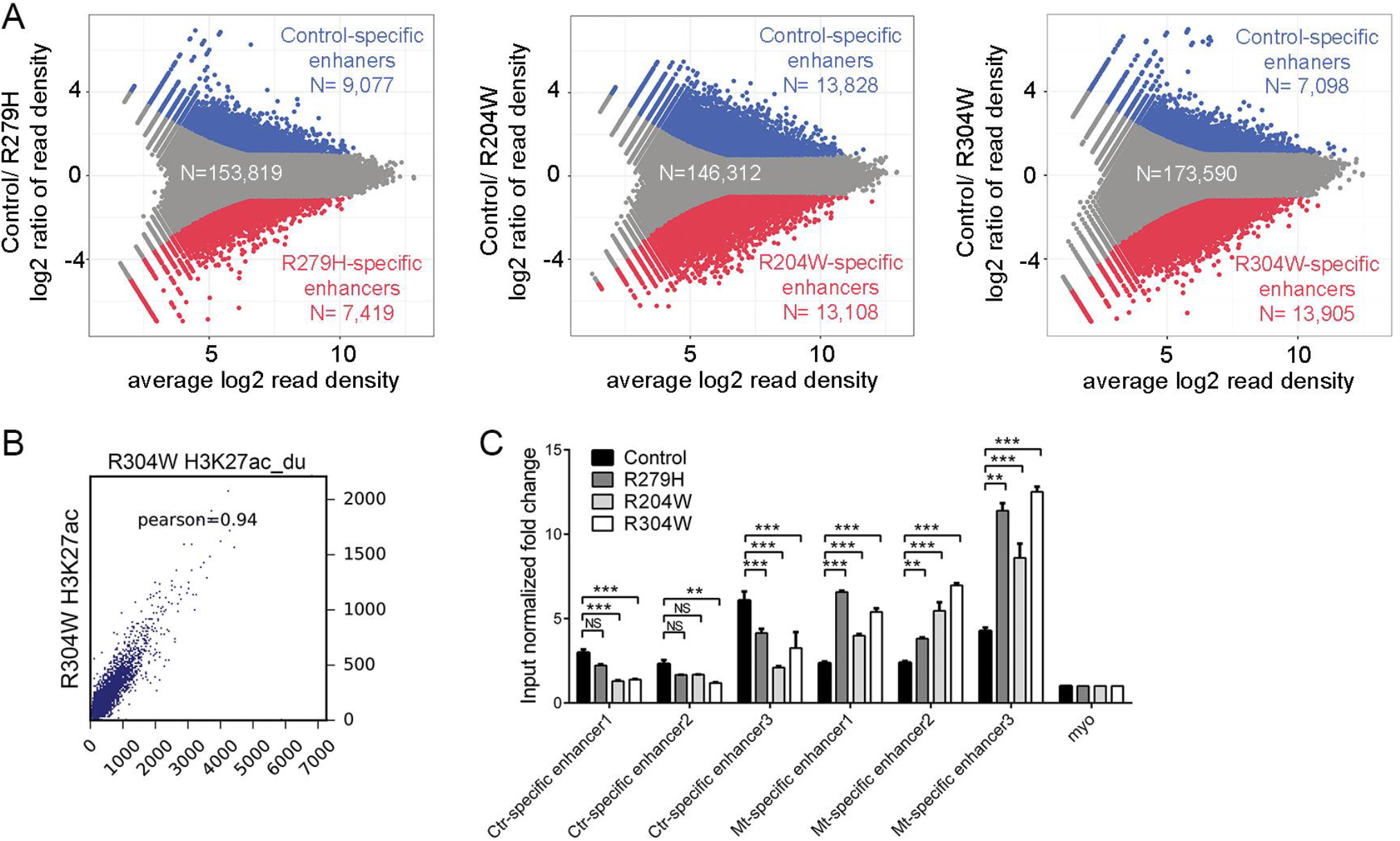
Transcriptional rewiring in p63 mutant keratinocytes, related to Figure 4. A. Differential H3K27ac signals in control and p63 mutant keratinocytes shown by the MAplots. Each dot represents an H3K27ac peak. Blue dots represent regions with significantly higher H3K27ac signals in control keratinocytes (Control-specific enhancers, *P* value <e10–5, M value >1); red dots represent regions with significantly higher H3K27ac signals in p63 mutant keratinocytes (R279H, R204W, and R304W-specific enhancers, from left to right, *P* value < 10e-5, M value < −1). Gray dots indicate non-differential peaks. B. Scatterplot showing correlation of signal intensity of H3K27ac peaks between the two replicas. Value displayed is the Pearson correlation coefficient. C. H3K27ac ChIP-qPCR validation of differential enhancers detected in the MAplots. Information about these three ‘control-specific’ enhancers and three ‘mutant-specific’ enhancers is detailed in Table EV1C. Input normalized fold change is relative to both input DNA and negative control loci (myo). Data are shown as mean ± standard deviation, *n*=2, two-way ANOVA, NS *P* value > 0.05, ** *P* value <0.01, *** *P* value <0.001.

**Figure EV6.**
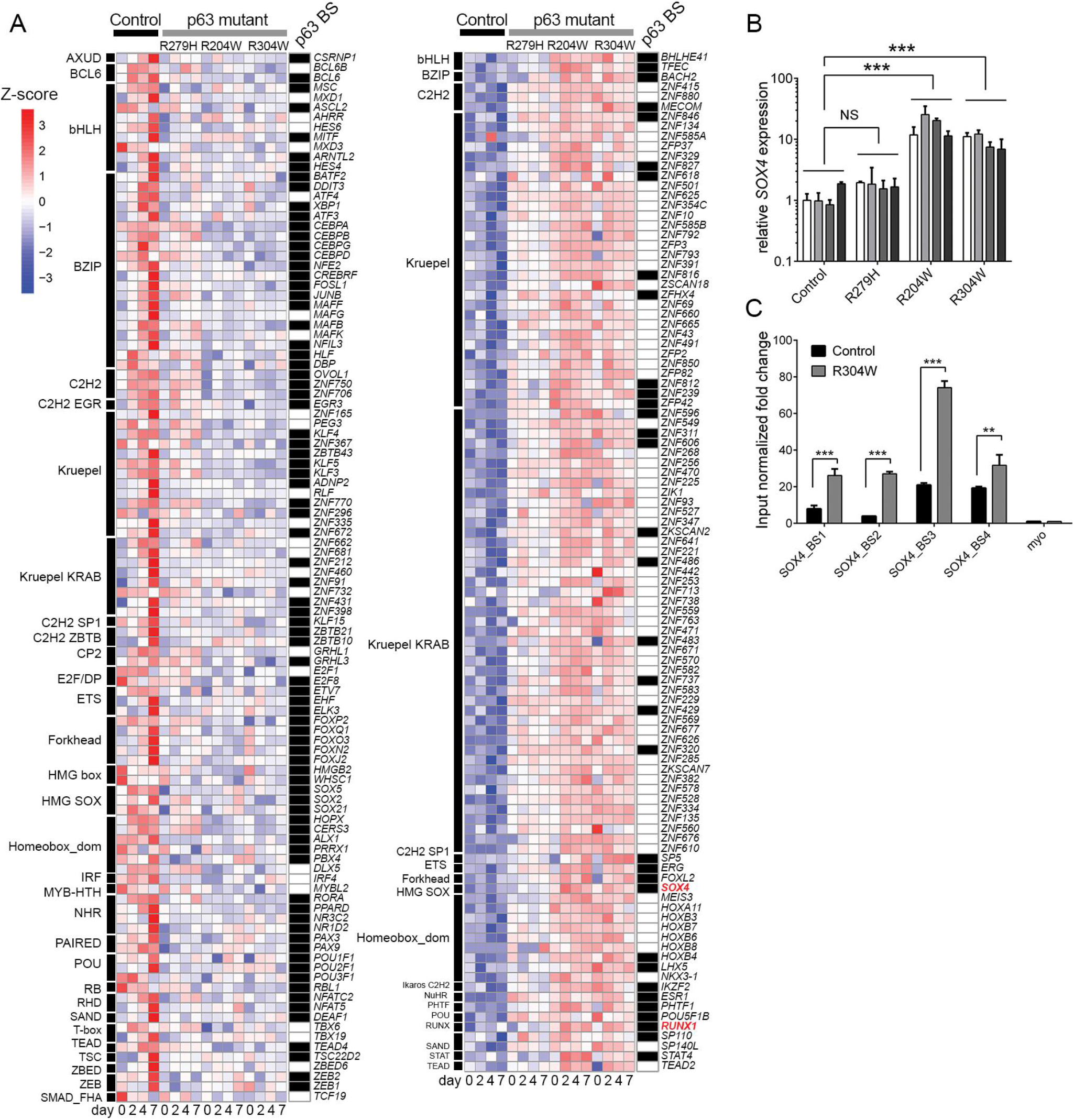
Deregulated TFs, related to Figure 4. A. Significantly downregulated (left) and upregulated (right) TFs in p63 mutant keratinocytes (*P* value <0.01 and fold change >2) shown by heatmaps. Nearby p63 BS of these genes are indicated, black, with at least one p63 BS; white, without p63 BS. TFs are grouped according to families. Z-score is calculated based on log10 (FPKM+0.01) to indicate the level of expression. B. Validation of *SOX4* gene expression analysis by RT-qPCR. Gene expression is normalized to the expression of *GusB* (reference gene). Primer information is provided in Table EV1C. Data are shown as mean ± standard deviation, *n*=2, two-way ANOVA, NS *P* value > 0.05, *** *P* value <0.001. C. SOX4 ChIP-qPCR performed in control and R304W mutant keratinocytes. Primer information about these SOX4 binding sites is detailed in Table EV1C. Input normalized fold change is relative to both input DNA and negative control loci (myo). Data are shown as mean ± standard deviation, *n*=2, two-way ANOVA, *** *P* value <0.001.

**Figure EV7.**
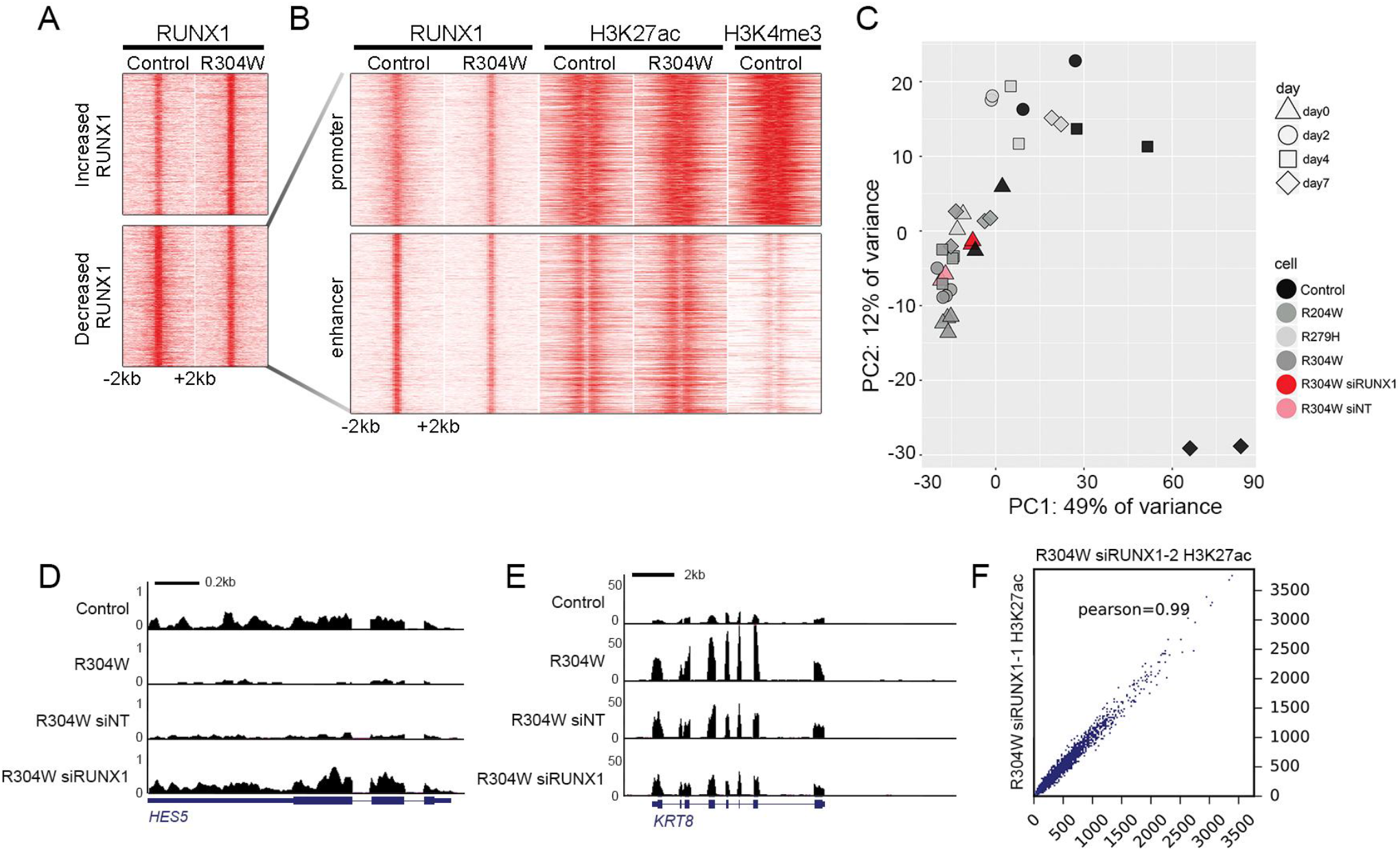
Reversing upregulated RUNX1 expression in p63 mutant keratinocytes can partially rescue deregulated gene expression and enhancer landscape, related to Figure 6. A. K-means clustering of merged RUNX1 binding sites combined from control keratinocytes and R304W mutant keratinocytes are shown in heatmaps in a 4 kb window with summits of RUNX1 in the middle (*k* = 2, metric = Pearson). Color intensity represents normalized read counts. B. Zoom-in re-clustering of decreased RUNX1 binding sites in R304W mutant keratinocytes. C. PCA plot on RNA-Seq data of RUNX1 knockdown in R304W mutant keratinocytes (R304W siRUNX1) and non-targeting siRNA treated keratinocytes (R304W siNT). The gene expression pattern of R304W siRUNX1 (red triangle) is closer to control keratinocytes (black triangle) than that of R304W siNT (pink triangle) and that of R304W mutant keratinocytes (grey triangle) at the proliferation condition. D. An example of rescued downregulated genes by RUNX1 knockdown. The UCSC genome browser screenshot of RNA-Seq tracks at gene loci of *HES5*. E. An example of rescued upregulated genes by RUNX1 knockdown. The UCSC genome browser screenshot of RNA-Seq tracks at gene loci of *KRT8*. F. Scatterplot showing correlation of signal intensity of H3K27ac peaks between the two biological replicas in R304W mutant keratinocytes treated with siRUNX1. Value displayed is the Pearson correlation coefficient.

## Expanded View Tables

Table EV1_Transcriptomic analysis of control keratinocytes and mutant keratinocytes during differentiation.xlsx

Table EV2_Chromatin landscape clustering during control keratinocytes differentiation.xlsx

Table EV3_ChromHMM based chromatin state characterization.xlsx

Table EV4_Characterization of p63 binding difference in control keratinocytes and p63 mutant keratinocytes.xlsx

Table EV5_Re-distribution of enhancers in p63 mutant keratinocytes.xlsx

Table EV6_Motif analysis of re-distributed enhancers in p63 mutant keratinocytes.xlsx

Table EV7_Characterization of RUNX1 binding difference in control keratinocytes and R304W mutant keratinocytes.xlsx

Table EV8_siRNA information.xlsx

Table EV9_Gene rescue by RUNX1 knockdown in R304W mutant keratinocytes.xlsx

Table EV10_Dataset summary.xlsx Table EV11_Western blots.xlsx

